# Biomechanical regulation of cell shapes promotes branching morphogenesis of the ureteric bud epithelium

**DOI:** 10.1101/2024.03.28.585666

**Authors:** Kristen Kurtzeborn, Vladislav Iaroshenko, Tomáš Zárybnický, Julia Koivula, Heidi Anttonen, Darren Brigdewater, Ramaswamy Krishnan, Ping Chen, Satu Kuure

**Affiliations:** Helsinki Institute of Life Science, University of Helsinki, Finland; Stem Cells and Metabolism Research Program Unit, Faculty of Medicine, University of Helsinki, Finland; Department of Biochemistry and Developmental Biology, Faculty of Medicine, University of Helsinki, Finland; Department of Pathology and Molecular Medicine, McMaster University, Hamilton, Ontario, Canada; Center for Vascular Biology Research, Department of Emergency Medicine, Beth Israel Deaconess Medical Center, Boston, MA, USA; Division of Clinical Chemistry, Department of Laboratory Medicine, Karolinska Institutet, Stockholm, Sweden; Laboratory Animal Centre, University of Helsinki, Finland

**Keywords:** branching morphogenesis, kidney development, mechanical forces, cell sizes, biomechanics

## Abstract

**Background:** Branching morphogenesis orchestrates organogenesis in many tissues including kidney, where ureteric bud branching determines kidney size and nephron number. Defects in branching morphogenesis result in congenital renal anomalies which manifest as deviations in size, function, and nephron number thus critically compromising the lifelong renal functional capacity established during development. Advances in the genetic and molecular understanding of ureteric bud branching regulation have proved insufficient to improve prognosis of congenital renal defects. Thus, we addressed mechanisms regulating three-dimensional (3D) ureteric bud epithelial cell morphology and cell shape changes during novel branch initiation to uncover the contributions of cellular mechanics on cellular functions and tissue organization in normal and branching-compromised bud tips.

**Methods:** We explored epithelial cell behavior at all scales by utilizing a combination of mouse genetics and a custom machine-learning segmentation pipeline in MATLAB. Ureteric bud epithelial cell shapes and sizes were quantified in 3D wholemount kidneys. A combination with live imaging of fluorescently labelled UB cells, traction force microscopy, and primary UB cells were used to determine how basic cellular features and niche biomechanics contribute to complex novel branch point determination in the process that aims at gaining optimal growth and epithelial density in a limited space.

**Results:** Machine learning-based segmentation of tip epithelia identified geometrical round-to-elliptical transformation as a key cell shape change facilitating shifts in growth direction that enable propitious branching complexity. Cell shape and molecular analyses in branching-compromised epithelia demonstrated a failure to condense cell size and conformation. Analysis of branching-compromised ureteric bud derived epithelial cells demonstrated disrupted E-CADHERIN and PAXILLIN mediated adhesive forces and defective cytoskeletal dynamics as detected by fluorescent labelling of actin in primary ureteric bud epithelial cells. Branching-compromised ureteric bud epithelial cells showed wrinkled nuclear shapes and alterations in MYH9-based microtubule organization, which suggest a stiff cellular niche with disturbed sensing of and response to biomechanical cues.

**Conclusions:** Our results indicate that the adhesive forces within the epithelium and towards the niche composed of nephron progenitors must dynamically fluctuate to allow complexity in arborization during new branch formation. The data collectively propose a model where epithelial cell crowding in tandem with stretching transforms individual cells into elliptical and elongated shapes. This creates local curvatures that drive new branch formation during the ampulla-to-asymmetric ampulla transition of ureteric bud.

## INTRODUCTION

Organogenesis of several different functional systems originating from distinct germ layers depends on complex epithelial tube morphogenesis commonly known as branching morphogenesis.^1–3^ Branching involves repeated formation of new branch points and tube elongation, which together facilitate the formation of a maximal functional area in a limited three-dimensional (3D) space. Two distinct mechanisms of branch point formation have been described: tip bifurcation (or clefting) and side (lateral) branching.^4, 5^ In lung and kidney, stereotyped branching morphogenesis dictates, but the patterns differ from each other by the presence or absence of lateral branching events. In the lung, three geometrically distinct modes of branching dictate: domain (lateral) branching, planar bifurcation, and orthogonal bifurcation, and each of these modes occurs in a fixed order.^6^ In the kidney, the ureteric bud (UB) undergoes a series of reiterative rounds of tip bifurcations early in development followed by some trifurcations during later embryonic development. The morphological changes in the UB tip give rise to an epithelial tree where the extent and gross pattern of branch elaboration are conserved and highly reproducible.^7, 8^

Branching morphogenesis of the epithelial UB is required to build a functional kidney.^9^ The significance of the branching process is highlighted by studies showing that mutations in genes regulating branching morphogenesis often lead to severe kidney malformations like aplasia, polycystic renal dysplasia, and hypoplasia.^10–13^ Reciprocal tissue interactions between the epithelial UB tip and the surrounding mesenchyme govern the branch pattern and are mediated by organ-specific secreted signaling molecules and extracellular matrix components.^1, 2^ It is well established that such reciprocal crosstalk is largely mediated by several growth factors expressed by the surrounding metanephric mesenchyme and that they typically activate receptor tyrosine kinase signaling in the UB.^14^ The most notable UB branching regulators include glial cell line-derived neurotrophic factor (GDNF) ^15–20^ and fibroblast growth factor (FGF) family members.^21–24^

Several intracellular cascades are activated when GDNF and FGFs bind to their receptor tyrosine kinase receptors on UB cell surfaces. Important for renal development are at least mitogen-activated protein kinase/extracellular signal-regulated kinase (MAPK/ERK), phosphoinositide 3-kinase/protein kinase B (PI3K/AKT), and phospholipase Cγ (PLCγ).^25^ These all regulate specific sets of transcriptional and cellular targets to control epithelial cell intrinsic and extrinsic events. Accumulating experimental evidence demonstrates that MAPK/ERK, which functions through the RAS-RAF-MEK-ERK cascade and activates many transcription factors and downstream protein kinases notably targeting proteins involved in cellular adhesion, including paxillin ^26^ as well as those involved with actin polymerization, such as the actin nucleator Arp2/3 complex ^27^, is essential for normal UB branch formation. This was originally suggested *in vitro* by chemical MEK inhibition studies in cultured kidneys ^28, 29^ and more recently demonstrated *in vivo* by genetic abrogation of the pathway specifically in the UB.^10^ Cellular characterization of MAPK/ERK-deficient UB epithelium identified abnormalities in focal adhesions and adherens junctions, while transcriptional profiling of these cells demonstrated loss of progenitor status leading to premature differentiation of collecting duct epithelium.^10, 30^

Extensive cell rearrangements and cell geometry changes sculpture epithelia in different developing organs across organisms.^31^ In essence, coordinated changes in the shapes of individual cells comprising the epithelium bend the tissue in a process leading to variations in morphological organization. Highly dynamic cellular adhesions are intimately coupled to tightly regulated cell shape changes, which contribute to tissue remodeling and morphogenesis, as described in many developing organs in *Drosophila*.^32–34^ It has also recently been shown that epithelial integrity and morphology in mature mammalian collecting ducts depend on the fine control of the actomyosin network dynamics involving actin polymerization, which was previously shown to be required for normal UB branching.^35, 36^ Inhibition of myosin function in mouse lung resulted in abnormal cell shapes suggesting functions in constraining cell morphology, which however must be released at the sites of new branch points.^37^ The function of two non-muscle myosin heavy chain proteins, MYH9 and-10, is essential for epithelial integrity and kidney UB.^38^ Furthermore, Rac1 controls the actin cytoskeleton to prevent misshapen cells from entering the cell cycle during renal tubule repair.^39^

In the developing mammalian kidney, the UB tip-localized progenitor cells are motile and undergo luminal mitosis ^40–42^ but the exact cell shape changes and their contribution to the stepwise process of UB branching morphogenesis are not characterized. Here we utilize a recently developed machine learning-based algorithm called ShapeMetrics ^43^ to systematically analyze cell shapes and their changes in normal and MAPK/ERK-deficient UB epithelium used as a model of branching deficiency. Our characterization of molecular and biomechanical consequences revealed that remodeling of cell adhesion molecules leads to changes in size, morphology, and traction stress of the cells within the UB tip niche of branching-compromised (BC) kidneys.

## METHODS

### I. Mice

All experiments were approved by the Finnish Animal Care and Use Committee (license KEK16-020) in compliance with all relevant ethical regulations regarding animal research. The design and genotyping of HoxB7CreGFP, *Mek1*-floxed, *Mek2*-null, beta-catenin-null and beta-catenin stabilized mice have been described.^24, 44–47^ Mice were of mixed genetic background with contributions from C57BL/6JOlaHsd. NMRI mice were used for MEK-inhibition experiments.

Kidneys were cultured in DMEM:F12 + GlutaMAX (Gibco) supplemented with 10% fetal bovine serum (FBS; Gibco), and penicillin-streptomycin (PS) (“full DMEM”) under 5% CO_2_ at 37°C.^48^ MEK-inhibition was achieved by the addition of 15 µM U0126 (Cell Signaling Technologies) in serum-free media (DMEM + PS) and controlled by addition of the same amount of dimethyl sulfoxide (DMSO) in serum-free media to control kidney cultures.

Mechanical separation of the ureteric buds from E11.5-12.5 kidneys was carried out according to published protocol.^49^ Briefly, enzymatic collagenase-treatment (4 mg/mL, 15 min at 37°C) was followed by collagenase neutralization with 25 u/mL DNase I in DMEM/F12 + 10% FBS. Isolated UBs were cultured on the fibronectin-coated coverslips in DMEM/F12 + 10% FBS medium supplemented with GDNF (5 ng/mL), FGF2 (25 ng/mL), and HGF (50 ng/mL).

### II. Cell lines and transfections

Cells from the UB-derived cell line (Barasch et al., 1996) were cultured in DMEM supplemented with 10% FBS, 1x GlutaMAX, and 1x Normocin. UB-dervied cells were transfected with green fluorescent protein (GFP)-tagged wildtype paxillin using Lipofectamine 3000 and co-stained with E-cadherin, endogenous paxillin, and overexpressed paxillin. The cell line was also transfected with GFP-tagged, dileucine- and dialanine-modified E-cadherin with improved surface expression.^50^ These cells were co-stained with E-cadherin, endogenous paxillin, and overexpressed E-cadherin.

### III. Immunohistochemistry, imaging, and machine learning-based segmentation

The isolated UBs and ureteric bud-derived cell line were fixed with 4% paraformaldehyde in PBS solution for 20 minutes at room temperature before being washed in PBS 3−10 minutes. The permeabilization has been performed with 0.1% Triton X-100 + 50mM glycine for 15 min, followed by PBS washes for 3−10 minutes with gentle rocking. Cells were blocked for 30 minutes with 3% FBS in PBS, followed by primary antibody staining overnight in a cold room. Samples were washed in PBS for 3−10 minutes before secondary antibody staining for 60 minutes. After 3−10 minute washes, nuclei were stained by Hoechst 33342 for 10 minutes. After follow-up PBS washes, cover slips were mounted using the Epredia™ Immu-Mount.

For whole-mount imaging, kidneys were dissected from embryos at E12.5, and shortly attached to Transwell filters (3h, Transwell permeable supports, 0.4 um polyester membrane, Corning Incorporated). For whole-mount imaging, samples were fixed for 10 min in ice-cold methanol and incubated with primary antibodies (goat E-cadherin, R&D systems) and nuclear stain (Hoechst) at 4°C twice o/n. Tissues were then incubated with secondary antibodies (anti-goat Alexa Fluor 488, Jackson Immuno Research laboratories) for 2 h at RT or o/n at 4°C. Stained kidneys were mounted with 99% glycerol (Sigma-Aldrich).

For paraffin sections, tissues were fixed with 4% paraformaldehyde (PFA), embedded in paraffin using an automatic tissue processor, and cut to 5 µm thickness. Paraffin sections were rehydrated by xylene-alcohol series. Antigen retrieval was performed by simmering sections for 5-15 minutes in Tris-HCl EDTA buffer. Sections were blocked with 10% FBS for 1 h at RT prior to o/n incubation with primary antibodies at RT or 4°C. Confocal micrographs of whole-mount and paraffin-sectioned samples were taken with Leica SP8 CARS. For cell segmentation with ShapeMetrics, pixel classification from 3D confocal images was performed using ilastik ^51^, and segmentation was performed with a custom algorithm in MATLAB (MathWorks). In brief, Ilastik creates prediction maps for MATLAB-based further processing of the image, which is followed by watershed segmentation and connection of possible gaps in cell borders. The segmented cells can be used to collate a single unbiased heatmap and divided into subgroups according to cell features that were used ^43^ In total, 11 BI and 3 BC UB tips were segmented (Table 1-2).

### IV. Extraction of spatial parameters and MATLAB analysis methods

The basic 3D reconstruction of UB tip cells, extractions of geometric parameters and clustergram analysis was done using original version of Shapemetrics script (https://github.com/KerosuoLab/Shapemetrics) as described previously.^43^ Additionally, a new script was written. This code sorts parameter values from highest to lowest and allows to select specific subgroups of cells for further characterization, as used here for the top 10% of cells. Geometric parameters that we extracted are volume, roundness, ellipticity, elongation, longest axis, minor axis, and intermediate axis. For visualization, we converted spatial data into numeric values for each parameter at a single cell resolution and performed principal component analysis and Uniform Manifold Approximation and Projection (UMAP) to two dimensions.

For both clustergram analysis and 10%-quantile method, merging of selected cells and original images was done in ImageJ. Merged multi pages were projected onto 2D images using standard deviation algorithm in ImageJ. Distribution and whisker plots were made from the matrices generated in Shapemetrics script by using OriginPro. Basic column statistics including average, median and skewness coefficients values were also calculated using OriginPro.

### V. Traction force microscopy

Traction force microscopy of cell doublets was conducted as described previously.^52, 53^ Micrographs were taken using a Zeiss Axiovert inverted microscope coupled to a CSUX1 spinning-disc device (Yokogawa) in an environmental chamber at 37°C and 5% CO_2._ In brief, polyacrylamide gels of 19.66 kPa elasticity were embedded with 0.2 µm 505/515 fluorescent beads (Life Technologies) and coated with fibronectin (R&D Systems). UB-derived cells were seeded at a density of 50,000 cells/gel to achieve cell doublet formation and incubated once o/n in full media and once more o/n with 15 µM U0126 or DMSO in serum-free media prior to imaging. After all doublets were imaged (experimental), 10X trypsin (Gibco) was added, and cells were completely detached. Gel fluorescent beads were imaged again in the absence of cells (reference). A custom Fourier-Transform Traction Cytometry algorithm was used to calculate tractions based on bead displacement between “experimental” and “reference” images, described in detail previously.^53^

For cell monolayers, a modified traction force microscopy approach was used.^54, 55^ Briefly, cells were seeded upon fibronectin-coated (R&D Systems) Softrac hydrogels (Matrigen) with 25 kPa stiffness, embedded with 0.2 um yellow-green fluorescent microspheres in 35 mm dishes. The experiment was performed in three parts: first with a cell line derived from renal UB epithelium at 80% confluence, then again at 100% confluence, and third with primary UB epithelium excised from embryonic kidneys and allowed to delaminate onto the gels. In each part, for each cell monolayer, we obtained fluorescent microspheres images with and without cells present. From the image pair, we determined the cell-exerted displacement field, and from the displacement field, the traction field using the method of Fourier-Transform Traction Cytometry, modified to the case of cell monolayers. We report traction as the root mean square value within the field of view (yy) and averaged them across cell monolayers from each experimental part. Images were taken using 3i Marianas attached to a fully-motorized Zeiss Axio Observer Z1 microscope at 37°C and 5% CO_2_.

### VI. On-Cell Western

Cells from the UB-derived cell line were plated at a density of 12,500 cells/well in a black, clear-bottom 96-well plate (Perkin Elmer). Prior to seeding, all wells were coated with 10 µg/ml fibronectin (R&D Systems). Cells were cultured in DMEM:F12, 10% FBS, GlutaMAX, and normocin under 5% CO_2_ at 37°C for 24-36 hours, until they reached a confluence of 80-90%.

After reaching the desired confluence, cells in 48 wells were treated with 15 µM U0126 in DMEM/normocin. Cells in the other 48 wells were controlled by addition of DMSO in DMEM/normocin. Cells were incubated with treatment for 24 h. Media containing U0126/DMSO was changed once during the incubation.

Cells in 60 wells were incubated with primary goat polyclonal anti-mouse E-cadherin (R&D Systems, Cat. No AF748, 1:200) in cold 5% FBS/PBS for 20 min on ice. Negative control cells (36 wells) were incubated only with cold 5% FBS/PBS. Cells were rinsed three times with cold PBS and fixed with 4% PFA for 10 minutes at room temperature. All wells were washed 3−5 mins with PBS. After washing, blocking was performed for 15 minutes with Odyssey blocking buffer (PBS) (LI-COR Biosciences, Cat. No 927-40000) at room temperature followed by 20 mins incubation in the dark with IRDye 800CW donkey anti-goat antibody (LI-COR Biosciences, Cat. No 926-32214, 1:1000) in 50% blocking buffer and 50% PBS/0.2% Tween-20. Cells were co-stained with nuclear stain: DRAQ5 (Thermo Fisher Scientific, Cat. No 62251, 1:7500) for normalization. Cells were then washed 3−10 mins with PBS/0.1% Tween-20. After washing, PBS was changed to wells and the plate was scanned with the LI-COR Odyssey IR Imager (169 μM resolution, 3 focus offset, and 6 to 8 intensity). Data was analyzed using Excel (Microsoft).

The mean intensity value of negative controls was subtracted from the 800/700 intensity ratio. Significance of variability between conditions was determined using the Student’s t-test (two-tailed, unpaired). Results with *P < 0.05, **P < 0.01, and ***P < 0.001 were considered significant. All experimental data are reported as means, and error bars represent experimental standard error (± standard deviation, SD).

### VII. Functional gene enrichment analysis

Functional gene enrichment analysis of differentially expressed genes identified in BC UB epithelium (HoxB7Cre-GFP;Mek1fl/fl;Mek2−/−) ^30^ was performed using the ToppFun application from ToppGene Suite (https://toppgene.cchmc.org) ^56^. GO enrichment of differentially expressed genes was performed with a false discovery rate of *p* < 0.05. GO biological processes shown in **Fig. S5** were selected based on their possible relation to the observed phenotype in branching-compromised UBs.

## RESULTS

Ureteric bud (UB) branching morphogenesis in the developing kidney utilizes repeated tip bifurcation followed by tube elongation to form a complex 3D network of epithelial tree (**Fig. 1**). During active branching morphogenesis the UB tip undergoes a series of well-described morphological changes.^7, 29^ The initial bud balloons to transform into an ampulla, which then grows and converts into an asymmetrical ampulla where the bifurcation event and changes in growth direction are determined (**Fig. 1A-C**). As a result, new bud invades into free space away from the parental duct. All these events occur through a multitude of transitory phases during which 3D UB morphology deviates from the basic forms in total of some 200 tips at embryonic day 14.5 (E14.5) kidneys (**Fig. 1D-F**). For simplicity, we divided UB tips into four categories based on their basic morphology: initial bud, ampulla, asymmetrical ampulla, and T-bud (bifurcated) tip (**Fig. 1G, S1**). This categorization is used throughout our analyses.

**Figure 1.**
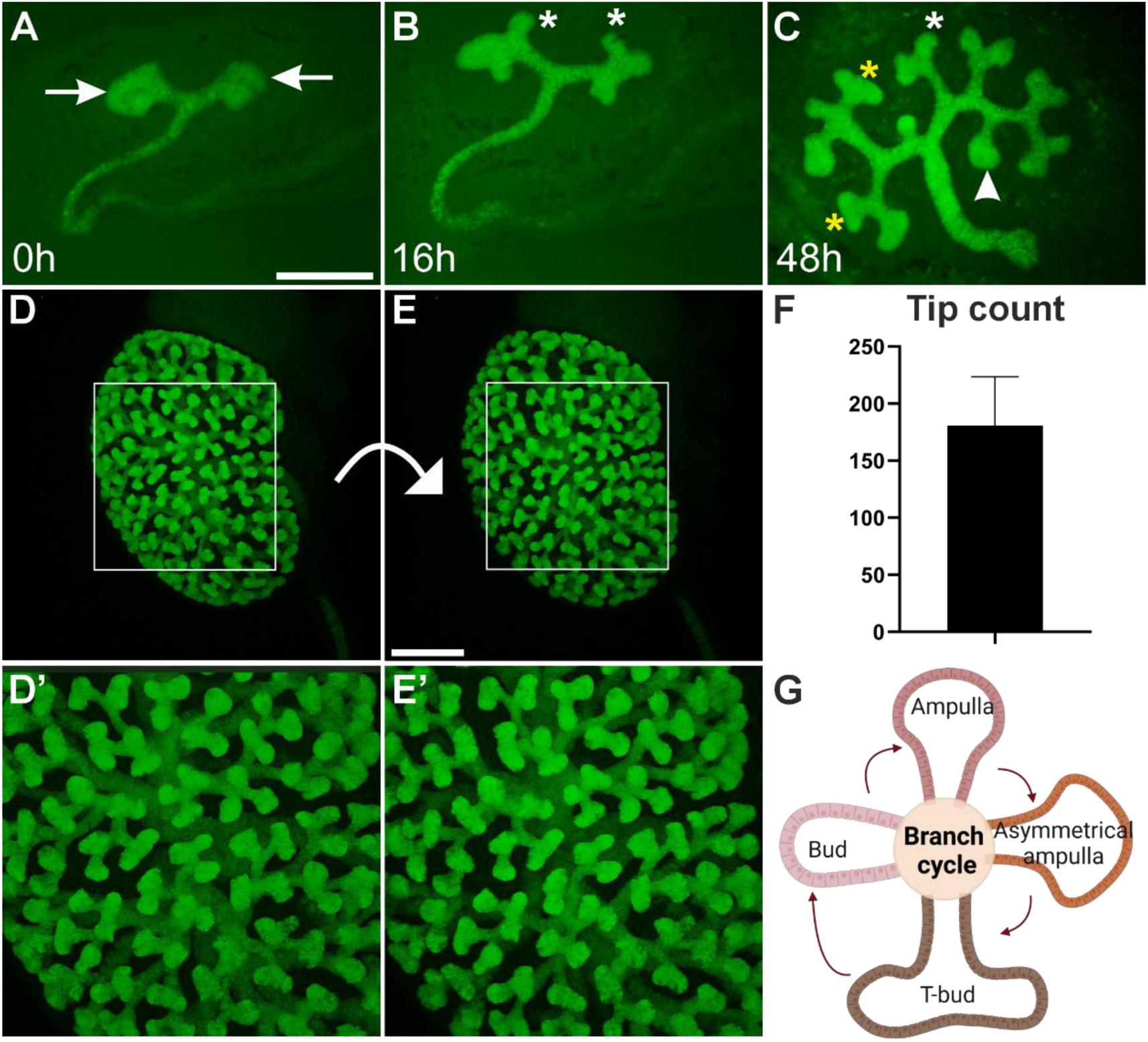
Ureteric bud branching morphogenesis in mouse embryonic kidney. E11.5 Hoxb7CreGFP kidney **A)** at the start (0h), **B)** 16h and **C)** 48h of *in vitro* culture. Green fluorescent protein (GFP) visualizes ureteric bud epithelium at its different morphological stages. White asterisk indicates a bud stage, arrowhead points to an ampulla stage, arrow highlights an asymmetric ampulla stage and yellow asterisks mark T-bud stage ureteric bud tip. **D-E)** E14.5 *HoxB7;Venus* kidney where ureteric bud tree is imaged in 3D and shown from different angles with appr. 35 degrees apart from each other. **D’-E’)** Zoomed in images of kidneys shown in D and E visualizing different ureteric bud tip morphologies in an intact kidney. **F)** Quantification of average ureteric bud tips in two kidneys at E14.5. **G)** Schematic presentation of repetitive ureteric bud branching cycle. Scale bar A-C: 500µm; D-E: 200µm

### Global quantification of cell shapes in branching-competent UB tips

To assess changes in UB cell sizes and shapes in an unbiased way, confocal z-stacks were obtained from entire, fixed kidneys at E12.5 immunostained with E-cadherin to mark epithelial membranes for the analysis of their specific geometrical features. We analyzed a total of 4030 epithelial cells originating from three different E12.5 mouse embryonic kidneys (**Table 1**). UB tips from the four distinct categories (**Fig. 1G; S1**) were quantified using basic geometrical parameters in the ShapeMetrics script: volume, roundness, elongation, ellipticity, and longest axis. Table 1 and S1 summarize the analyses and the median parameter values for segmented UB cells in each tip type.

The ShapeMetrics pipeline generates a hierarchical clustergram of cell geometries in segmented cells thus grouping epithelia based on their similarities and differences. Geometrical parameter values from the selected UB epithelial tips were concatenated into a joint matrix using hierarchical clustering in MATLAB. The clustergram demonstrates noticeable variety in cell shapes, thus demonstrating highly heterogeneous cell populations in the UB tips during branching (**Fig. 2A**). The ability to map cells with selected features back onto the original 3D biological sample provides spatial information about cell geometries in their native environment. By doing so we observed that pseudo-colored epithelial cells originating from the different clusters show high congestion and least variation in their localization through ampulla-to-T-bud stages (**Fig. 2B-E**).

**Figure 2.**
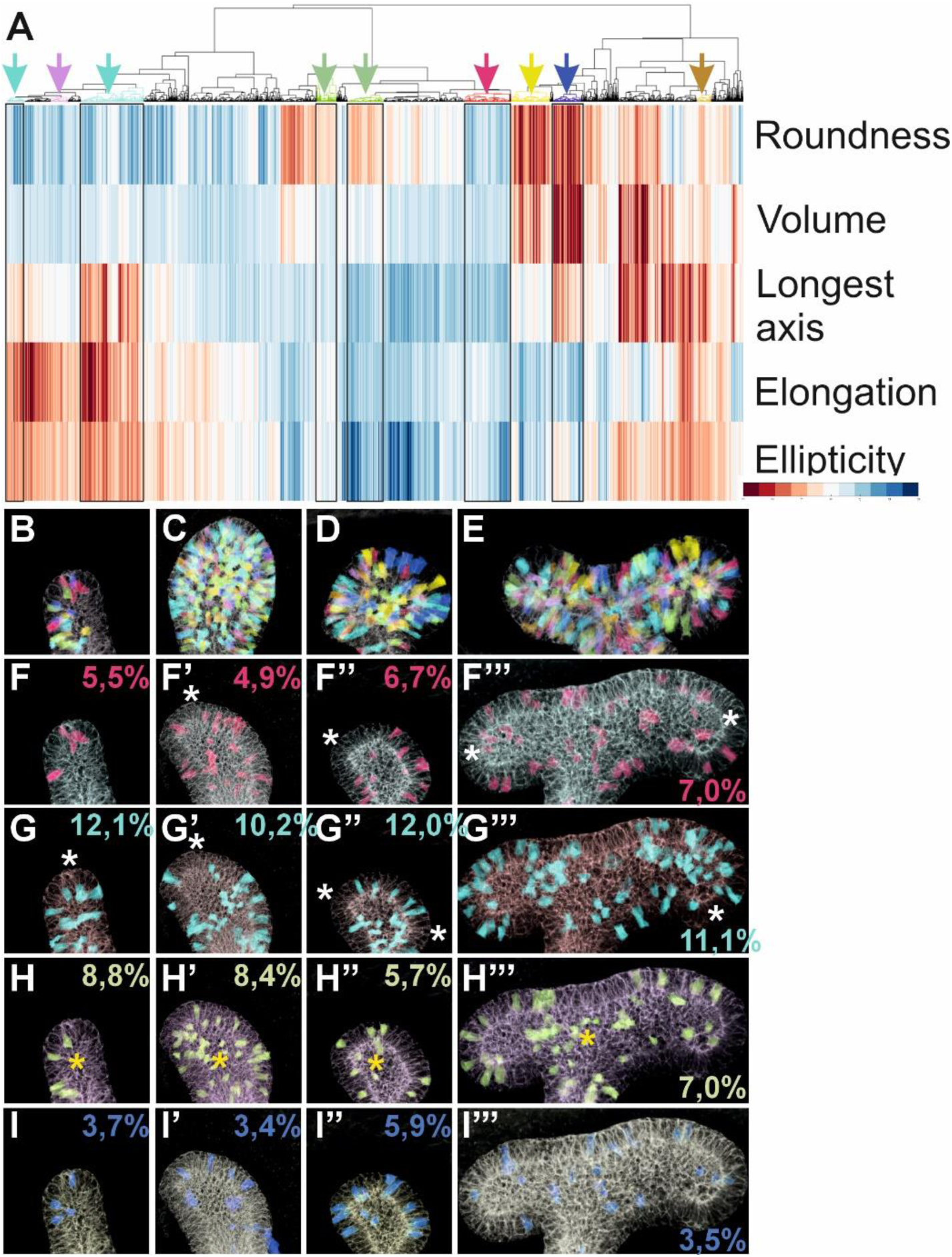
Heterogeneity in cell shape across the different stages of branch cycle tips. **A)** ShapeMetrics supervised machine-learning based algorithm was used to generate a heatmap cluster gram of 4030 segmented ureteric bud cells in three different E12.5 mouse kidneys. Each column represents a single cell, the color indicates the measured value of given geometrical parameter (blue: low; red: high), which are listed in right (roundness, volume, longest axis, elongation, ellipticity). Boxed areas highlight selected cell clusters, which are mapped back with corresponding colors to original biological samples of **B)** tip, **C)** ampulla, **D)** asymmetric ampulla, and **E)** T-bud stage ureteric bud epithelium. Cells in the clusters indicated by black rectangles are mapped back original samples at different branch cycle stages. Red cluster: low values for all geometrical parameters mapped back in the **F**) tip, **F’)** ampulla, **F’’)** asymmetric ampulla, and **F’’’)** T-mapped back to **G**) tip, **G’)** ampulla, **G’’)** asymmetric ampulla, and **G’’’)** T-bud stage ureteric bud epithelium. White asterisks indicate the most cortical UB tip regions. Green cluster: mapping back cells with high roundness to **H**) tip, **H’)** ampulla, **H’’)** asymmetric ampulla, and **H’’’)** T-bud stage ureteric bud epithelium. Yellow asterisk deciphers lumen of the epithelium. Blue cluster: mapping back epithelial cells with high roundness, volume, and longest axis to **I)** tip, **I’)** ampulla, I**’’)** asymmetric ampulla, and I**’’’)** T-bud stage ureteric bud epithelium. Numbers in figures F-I show the average proportion of cells that were quantified in each cluster from all biological samples analyzed. Scale bar: 40µm

### Highly heterogenic ureteric bud cells undergoing specific morphological changes during branching morphogenesis

Next, cells in clusters representing low volumes for all analyzed parameters and those showing increasing degree of complexity in combinations of elongation, ellipticity, volume and roundness were distinctly mapped back to the original UB tips. This revealed separate localization patterns for cells in all analyzed clusters (**Fig. 2F-I**) demonstrating preciseness of the ShapeMetrics tool. Cells with low values for all parameters were present with highest abundance of all segmented cells at T-bud stage (7,0%) and showed little variation in their localization patterns across different stages of entire branch cycle (**Fig. 2F, Table S2**). Cells that are of high elongation, ellipticity and longest axis predominated in abundance at initial bud and asymmetric ampulla stages (12,1% and 12%, respectively) and were more abundant across the entire branch cycle than cells in any other cluster (**Fig. 2G**). The round cells showed lowest abundance at asymmetric ampulla stage (5,7%) and higher luminal localization at ampulla and T-bud stages than at initial and T-bud stages (**Fig. 2H**). This could reflect at least two important biological features of the developing kidney: proliferation and spatial constrictions. Previous studies have shown that proliferation in the UB epithelium occurs at least partially through luminal mitosis, and epithelia in general are composed of confined space, where increase in cell numbers generates crowding phenomenon ^57, 58^. The observations that the cells with simultaneously high roundness, large volume, and high longest axis were the least abundant than any other cell cluster at all other but asymmetric ampulla stage shows special uniqueness for such geometrical combination in UB epithelium (**Fig. 2I**). Their paucity especially at highly proliferative ampulla stage ^15^ in relationship with cells that are of high roundness only (**Fig. 2H**) or those with highly elongated and elliptical form (**Fig. 2G**) demonstrates that smaller and columnar cells fit better into a limited space of epithelial sheet.

### Epithelial cells with the highest roundness peak at asymmetric ampulla stage while elongation increases steadily until the start of new branch cycle

Next, overall epithelial cell features were quantitated at different UB branch cycle stages. This uncovered that roundness is a parameter that fluctuates the most between the different branch cycle stages (**Table S1**). Comparison across the different branch cycle stages demonstrated that epithelial cell roundness is rather high already at the bud stage, while the ampulla stage shows diminished roundness, which then peaks at the asymmetric ampulla (**Fig. 3A**). This is supported by the distribution analysis, which show that difference in roundness is highest between asymmetrical ampulla and bifurcated tip stages, displaying the smallest overlap of roundness values (**Fig. S2A**) and suggesting major morphological changes in cell shapes during this transition. Interestingly, changes in cell roundness do not correlate with cell volumes, which remain virtually unchanged throughout the whole branch cycle progression (**Fig. 3B**), demonstrating that changes in cell roundness are not due to cells simply modifying their volume. Although average cell volume is similar at all UB tip stages (**Table 1**: 80µm^3^, 101 µm^3^, 91 µm^3^ and 100 µm^3^) the ampulla and T-bud tip stages occupy a wider range of volumes (45-485 µm^3^ and 65-500 µm^3^) including also higher volume cells that are not present at asymmetrical ampulla and initial bud stages (**Fig. S2B, E**). This indicates greater variation in cell sizes at ampulla and T-bud stages than at other UB morphologic stages.

**Figure 3.**
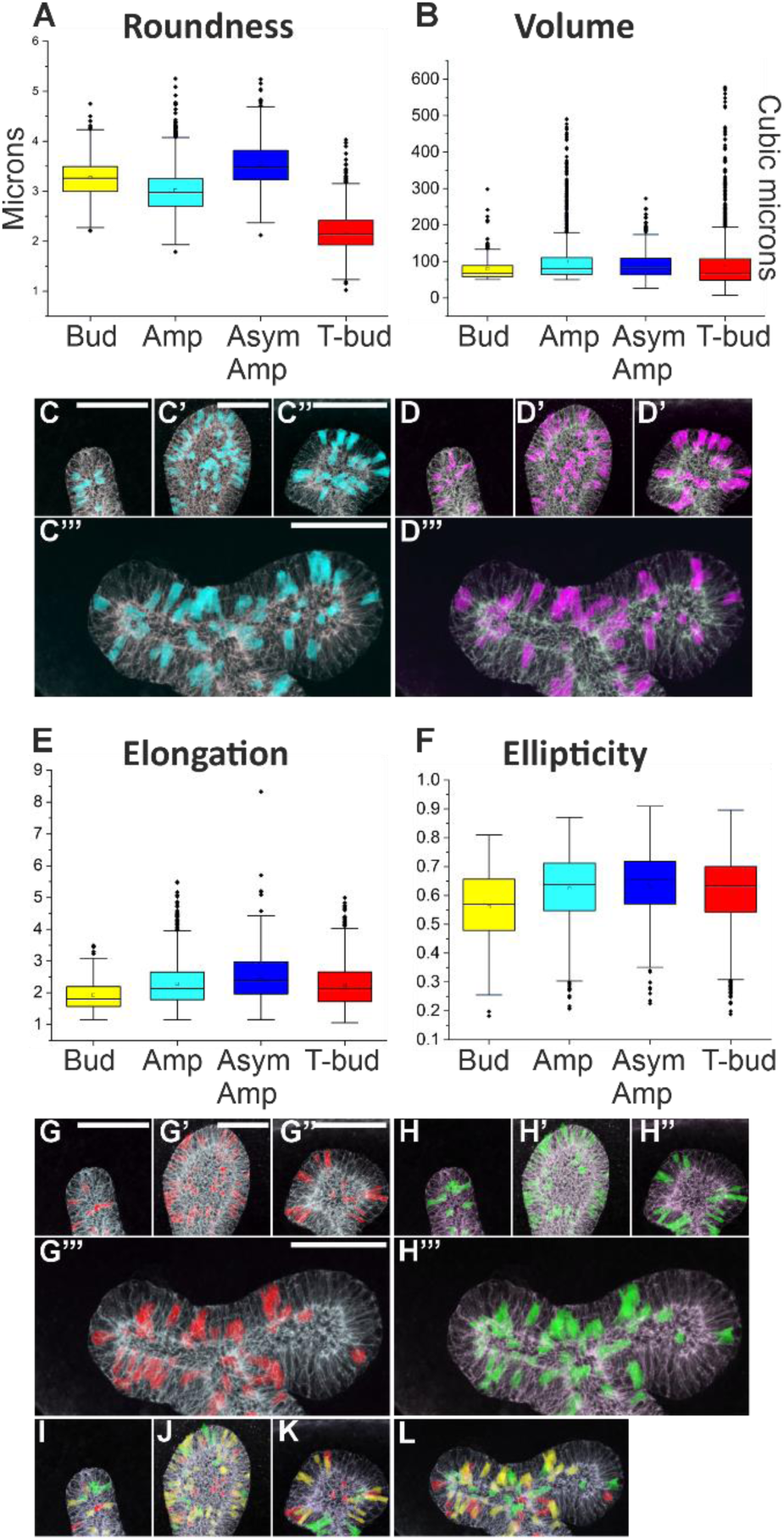
Cells with highest roundness fluctuate between different branch cycle stages while ellipticity increases until asymmetric ampulla. **A)** Quantitation of epithelial cell roundness across different stages of branch cycle demonstrates that asymmetric ampulla houses the most round ureteric bud cells. **B)** Quantitation of epithelial cell volume across different stages of branch cycle shows that cellular volume fluctuates very little between different tip morphologies. Top 10% of the most round epithelial cells (turquoise) mapped back to the original biological sample at **C)** bud, **C’)** ampulla, **C’’** asymmetric ampulla and **C’’**’) T-bud stage Top 10% of the high-volume epithelial cells (violet) mapped back to the original biological sample at **D)** bud, **D’)** ampulla, **D’’)** asymmetric ampulla and **D’’**’) T-bud stage. **E)** Quantitation of epithelial cell elongation across different stages of ureteric bud branch cycle. **F)** Quantitation of epithelial cell ellipticity across different stages of ureteric bud branch cycle. Top 10% of the most elongated epithelial cells (red) mapped back to the original biological sample at **G)** bud, **G’)** ampulla, **G’’** asymmetric ampulla and **G’’**’) T-bud stage. Top 10% of the most elliptical epithelial cells (green) mapped back at **H)** bud, H**’)** ampulla, H**’’)** asymmetric ampulla and H**’’’)** T-bud stage. Rainbow overlay of top 10% most elongated (red) and elliptical (green) cells at **I)** bud, **J)** ampulla, **K)** asymmetric ampulla and **L)** T-bud stage of ureteric bud demonstrates that very few cells with highest values for elongation and ellipticity are same cells. Abbreviations: Bud; initial bud, Amp; ampulla stage, As amp; asymmetric ampulla, T-bud; t-bud stage. Scale bar: 40µm

To characterize where the roundest, most elongated, most elliptical, and highest volume cells localize in branching UB tip, the top 10% cells of each parameter were mapped back to the original images. The roundest and highest volume cells showed similar overall pattern of localization in the UB epithelium across the different tip morphologies (**Fig. 3C-D**). Elongation, which is a parameter calculated as *<u>longest axis/average of intermediate</u> <u>and minor axes</u>*, gradually increases during the branch cycle progression reaching its peak at asymmetrical ampulla stage to decrease again at T-bud stage (**Fig. 3E, S2C**). In other words, cells elongate the most just before the bifurcation event. On the other hand, T-bud tip epithelia have the highest average lengths in all three dimensions: longest axis, intermediate axis, and minor axis (**Fig. S2G-I**). Changes in the epithelial cell ellipticity repeat the pattern seen for elongation while localizing sparsely in the initial bud, occupy ampulla and asymmetric ampulla unevenly, and are mostly devoid from the uttermost tip regions at T-bud stage (**Fig. 3F-H, S2C-D, F**). The most elliptical cells spatially differentiate from the most elongated cells, mainly by showing a distinct identity and localization when simultaneously mapped back to the biological sample (**Fig. 3I-L, S2C-D**).

### UMAP clustering of UB tip cells demonstrated T-bud stage singularity

We also carried out single-cell Uniform Manifold Approximation and Projection (UMAP) based on geometrical parameters to infer the developmental trajectory of epithelial cell shapes in UB tips (**Fig. 4**). This demonstrates dynamic changes in elongation throughout UB tip stages by showing noticeably close clustering of cell geometries at initial bud, ampulla, and asymmetrical ampulla stages. The cells at initial bud stage localize to the UMAP regions showing low elongation, at ampulla stage they appear to overlap with both initial bud and asymmetrical ampulla clusters. At asymmetrical ampulla stage cell geometries are shifted to the left representing the most elongated, elliptical, and round cells. The cells at T-bud tips cluster noticeably distinctly from the three other UB tip types showing low roundness with similar patterns of elongation and longest axis.

**Figure 4.**
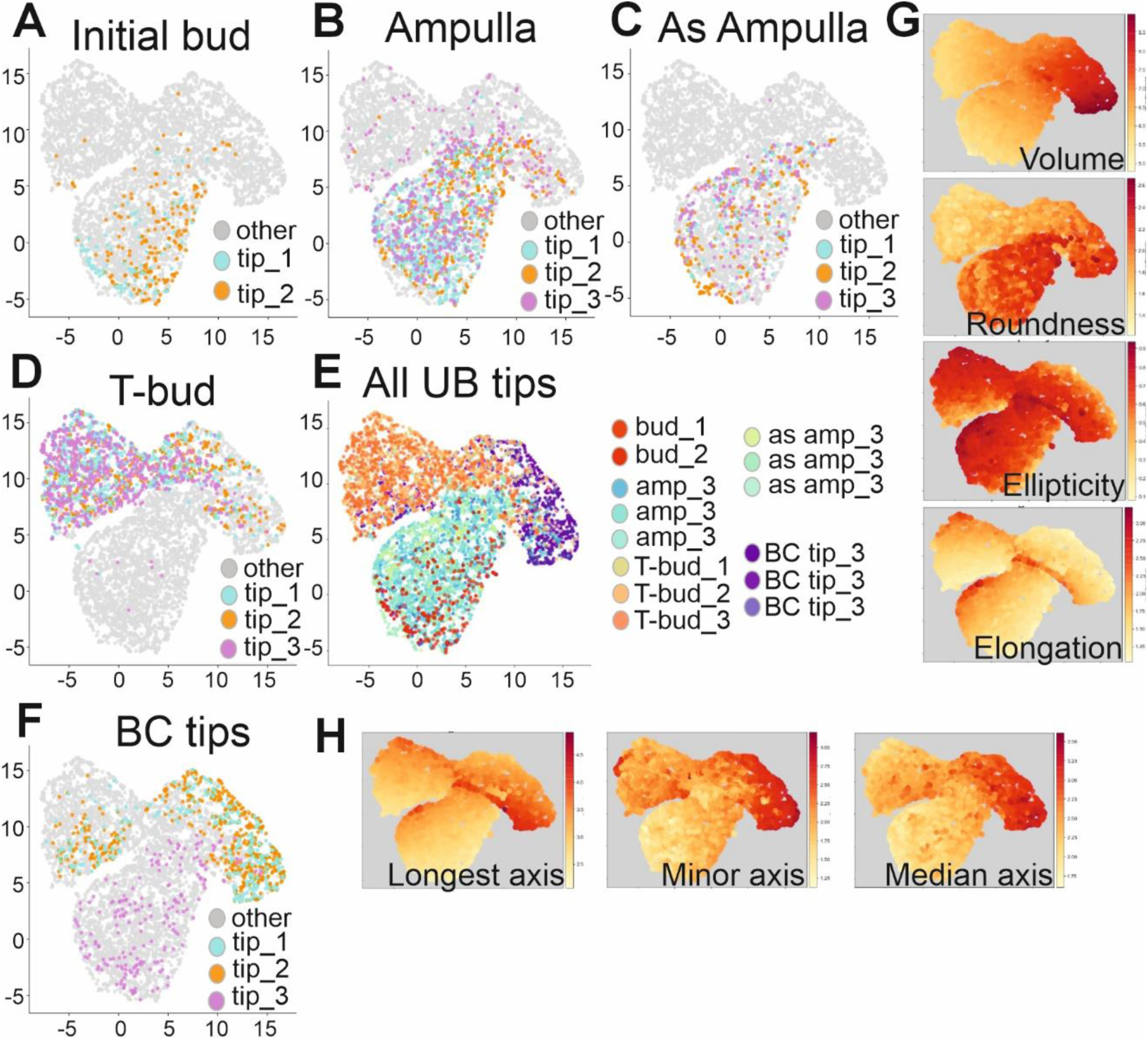
UMAP projection of ureteric bud epithelial cell reveals the individual nature of T-bud stage. UMAP projection of cells segmented at **A)** initial bud stage, **B)** ampulla stage, **C)** asymmetrical ampulla stage, **D)** T-bud stage, **E)** all UB stages including BI tip, and **F)** UMAP projection of cells segmented in branching compromised (BC) tips only. **G)** and **H)** UMAP visualization colored by the levels of each parameter; intense red = high value, light yellow = low value, intense red = high value, light yellow = low value.

### Cell shape changes in branching-compromised UB epithelium

To understand the relationship between cell shapes, geometry and branching we proceeded to study a branching-compromised (BC) UB epithelium lacking MAPK/ERK activity (*HoxB7CreGFP;Mek1^fl/fl^;Mek2^-/-^*) and displaying a simplified arborization phenotype.^10^ UMAP trajectory suggest that the overall cell features in two out of three BC tips cluster closest to T-bud tips of branching-intact (BI) epithelium, with the exception that they do not overlap in the regions of high elongation and ellipticity (**Fig. 4E-F**). *In vitro* time-lapse analysis of the UB tip morphologies during branch cycle progression revealed that most of the BC tips morphologically resembled the initial bud tip stage, while few deflections seemed to progress towards abnormal ampullae and asymmetric ampullae morphologies (**Fig. 5A-F**). Since the new UB growth directions are determined mainly in the tips,^3^, ^5^ the simplified UB morphology indicates failure to branch.

**Figure 5.**
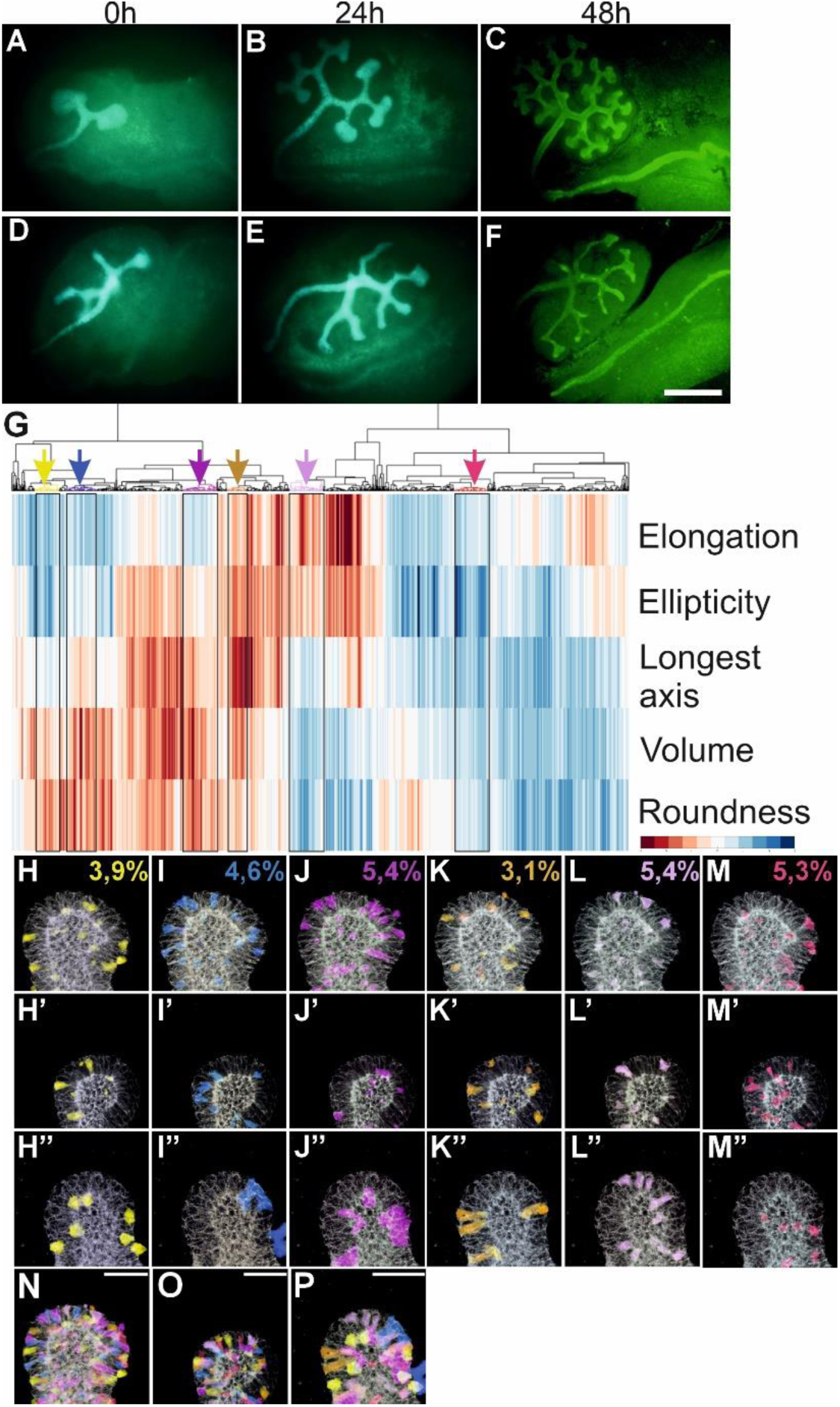
Overview of epithelial cell segmentation in branching-compromised ureteric bud. **A-C)** Time-lapse imaging of *in vitro* cultured branching-intact (BI, control) embryonic kidneys show full spectrum of ureteric bud morphological changes as visualized by GFP signal (green) during live imaging (A-B) and by CALBINDIN immunofluorescent staining (green) at 48h. **D-F)** Time-lapse imaging of *in vitro* cultured branching-compromised (*HoxB7CreGFP;Mek1^fl/fl^;Mek2^-/-^*) demonstrates less complex ureteric bud tip structures and impaired growth. **G)** ShapeMetrics algorithm was used to generate a heatmap cluster gram of 958 segmented ureteric bud cells in three different branching-compromised E12.5 mouse kidneys. Cells are clustered according to their distinct features, and each vertical column represents a single cell. The color of the column indicates the measured value of given geometrical parameter (blue: low; red: high), which are listed in right blue, violet, brown, lila, red) highlight the selected cell clusters that are mapped back with corresponding colors to original biological samples in H-M’’. **H-H’’**) Yellow cluster contains cells that are round and large in volume, **I-I’’)** blue cluster is otherwise similar but has higher longest axis, **J-J’’**) purple cluster cells have rather high values for all other parameters but elongation, **K-K’’**) brown cluster cells show high values for all parameters **L-L’’)** lilac cluster contains cells that are elongated and elliptical, and **M-M’’)** red cluster cells that show low values for all parameters. Numbers in figures H-M show the average proportion of cells that were quantified in each cluster from all biological samples analyzed. **N-P)** Images showing physical localization of all the selected cell clusters with the corresponding colors in the original biological samples. Scale bar: A-F: 500µm; H-M’’: 40µm.

Three BC tips with immature ampulla morphologies (**Fig. S3**) were subjected to ShapeMetrics analysis. Like in BI UB tips, heatmap clustering demonstrated heterogeneity of epithelial cell shapes in BC tips (**Fig. 5G)**. The BC cells from different clusters representing cells with high or low values for all parameters and those with increasing complexity in combinations of elongation, ellipticity, volume and roundness were mapped back to original samples (**Fig. 5H-M**). BC tip cells with high volume and roundness localized sparsely and were mainly found in the basal side of the epithelium showing variation in the intraepithelial localizations (**Fig. 5H**). Interestingly, similar cells which additionally are high in longest axis were showed increased abundance when compared to similar cells with shorter longest axes (**Fig. 5H-I**, 4,6% vs. 3,9%, respectively). Interestingly, BC cells that are of high roundness, volume and longest axis were more abundantly represented among all segmented cells in BC epithelia than similar cells at initial bud (3,7%) and ampulla stages (3,4%) of BI epithelia (**Fig. 2I-I’**). In BC epithelia the cells with high values for all other parameters but elongation and those with high elongation and ellipticity represented equal share of all segmented cells (5,4%), while the cells where all parameters are high were less abundant (3,1%) (**Fig. 5J-L**). Epithelial BC cells with low values for all parameters were present with similar abundance as in BI epithelia (5,3% vs. 4,9-7,0%, respectively) and predominantly localized to the trunk of the tips in 2/3 kidneys, while the tip cells in the one kidney cells show no localization preference (**Fig. 5M**). Interestingly, the overlay of all six subclustered epithelial cell groups demonstrated lose spatial localization and thus higher segregation of geometrical parameters in the BC than BI UB epithelium (**Fig. 5N-P**).

### Branching-compromised ureteric bud epithelial cells are high in volume and fail to compress into elliptical form

Our analyses of normally branching epithelium identified dynamic changes in cell roundness during the branch cycle progression as well as a steady increase of ellipticity and elongation up to the asymmetrical ampulla stage (**Fig. 3-4**). Since the BC UB epithelium demonstrates clearly less complex tip morphologies, we next compared general cell geometries between the two types of epithelia. This global quantification of cell shapes identified cell volume as one of the major differences between BI and BC UB tip cells (**Table1-2, S1**). Remarkably, the cellular volumes in BC tips are more than 2.5 times higher than epithelial cell volumes in the BI tips. The other parameters such as ellipticity and elongation in BC epithelial (0.54 and 1.93) are similar or lower than those detected in BI epithelium at the initial bud stage (0.57 and 1.98, **Tables 1-2**), supporting the morphological observation that BC epithelia are stuck at the bud stage of branch cycle. However, when cell roundness is used to characterize BC epithelial cells (3.07), they most closely resemble BI cells at the ampulla stage (2.97, **Tables 1-2**).

Mapping back the top 10% of the highest volume and roundest epithelial cells to BC epithelia shows that most of the high-volume BC cells are not the roundest cells (**Fig. 6A-D**). This suggests that other geometrical conformations also contribute to the high-volume cells. Indeed, some of the top 10% elliptical and elongated cells have large volumes, and thus likely compose a certain proportion of the highest volume cells in BC epithelium. Importantly, comparison of the BC epithelial parameters to those in BI epithelia across the entire branch cycle highlights differences in cell volumes but also identifies dynamic ascending variation in roundness, ellipticity, and elongation (**Fig. 6F-G**, **Fig. S4A**).

**Figure 6.**
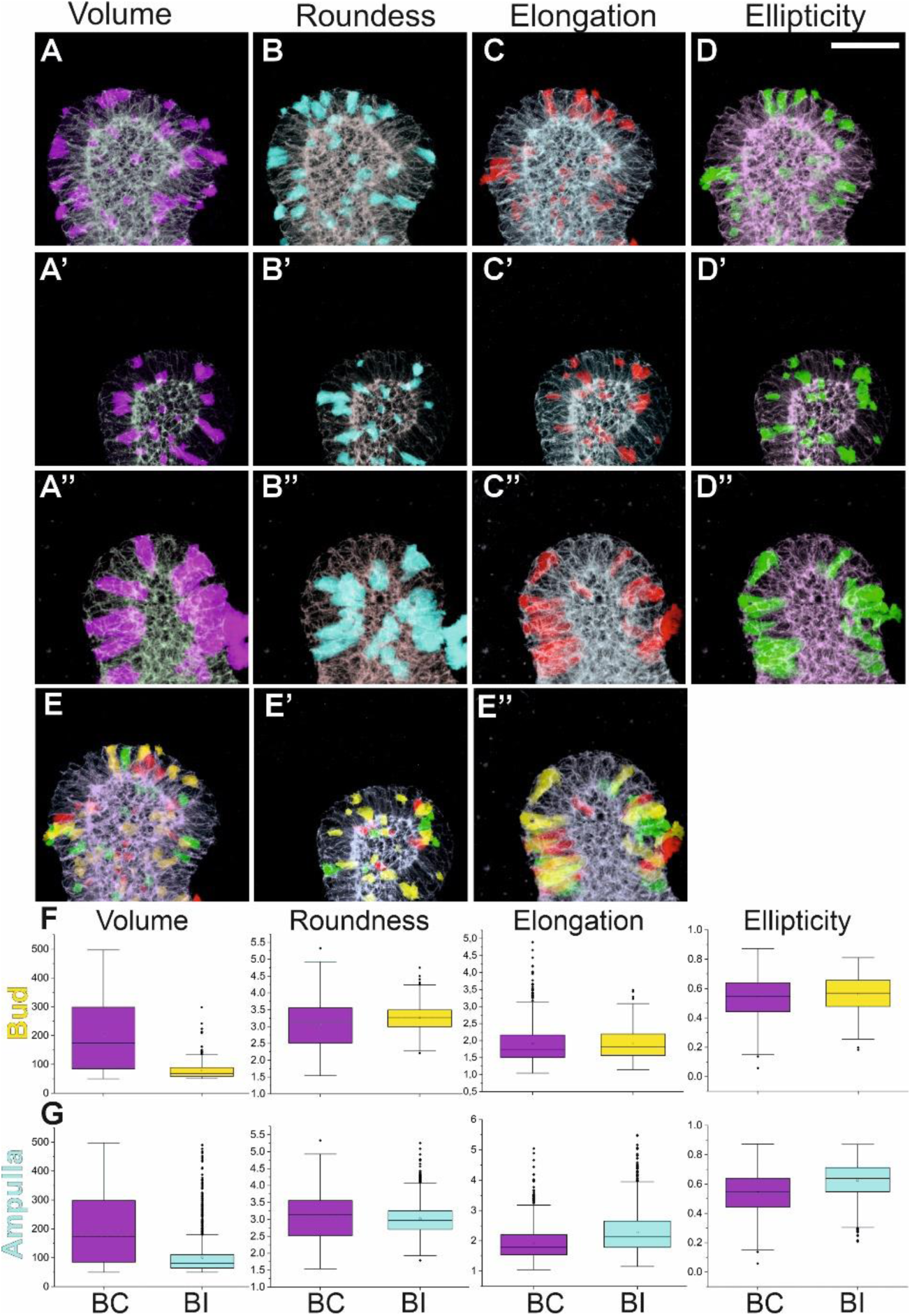
Branching-compromised ureteric bud epithelial cells are high in volume. **A-A’’)** Top 10% of the high-volume epithelial cells (purple, µ^3^) mapped back to three distinct branching-compromised ureteric bud. **B-B’’)** Top 10% of the most round epithelial cells (turquoise) mapped back to the three biological replicates of branching-compromised epithelium. **C-C’’)** Top 10% of the most elongated epithelial cells (red) mapped back to the three biological replicates of branching-compromised epithelium. **D-D’’)** Top 10% of the most elliptical epithelial cells (green) mapped back to the three biological replicates of branching-compromised epithelium. **E-E’’)** Overlay of the top 10% most elongated and elliptical cells localization in the three biological replicates of branching-compromised epithelium. **F)** Comparison of epithelial cell geometries in branching-intact (BI, yellow) and -compromised (BC, violet) ureteric bud at initial bud stage. Whisker plots showing volumes, roundness, elongation and ellipticity in epithelial cells of BI (violet bars) and BC (yellow) ureteric bud demonstrate marked variation in epithelial cell volume, which is significantly bigger in BI ureteric bud tips. **G)** Comparison of epithelial cell geometries in branching-intact (BI, yellow) and - compromised (BC, violet) ureteric bud at ampulla stage. Similarly as at earliest stage of ureteric bud branch cycle, epithelial cell volume remains higher in BC than BI ureteric bud tips. Additionally, epithelial cell roundness, elongation and ellipticity values are smaller in BC ureteric buds (violet bars) than those in BN (blue) buds. Abbreviations: BC; branching-compromised, BI; branching-intact. Scale bar: 40µm

The maximum cell volumes at initial bud (297µm^3^) and asymmetrical ampulla stages (272µm^3^) of BI epithelia were significantly lower than in BC tips (497µm^3^). In contrast, the volume range was comparable between the epithelial cells in BC tips and BI ampulla/T-bud stage tips (50-500µm^3^). To avoid bias arising from volume range differences for initial bud and asymmetrical ampulla stages, the comparisons to BC tips were done within the common cell volume range of 50-297µm^3^ for initial bud and 50-272µm^3^ for asymmetric ampulla stage (**Fig. 7**). This showed that cell volume range in BC tips is remarkably wider than in BI at initial bud and ampulla stages but shows better sharing at asymmetric and T-bud stages (**Fig. 7A-B, S4**). The distribution of the roundness parameter in BC tips almost completely overlaps with roundness distribution at BI ampulla stage, while the roundness distribution is slightly narrower in BI bud stage epithelial cells (**Fig. 7C-D)**. In contrast, at the asymmetrical ampulla stage, BI epithelial cells display generally higher roundness than cells in BC tips, and the situation is reversed when comparison is done between BC and BI epithelia at T-bud stage (**Fig. S4A-B**).

**Figure 7.**
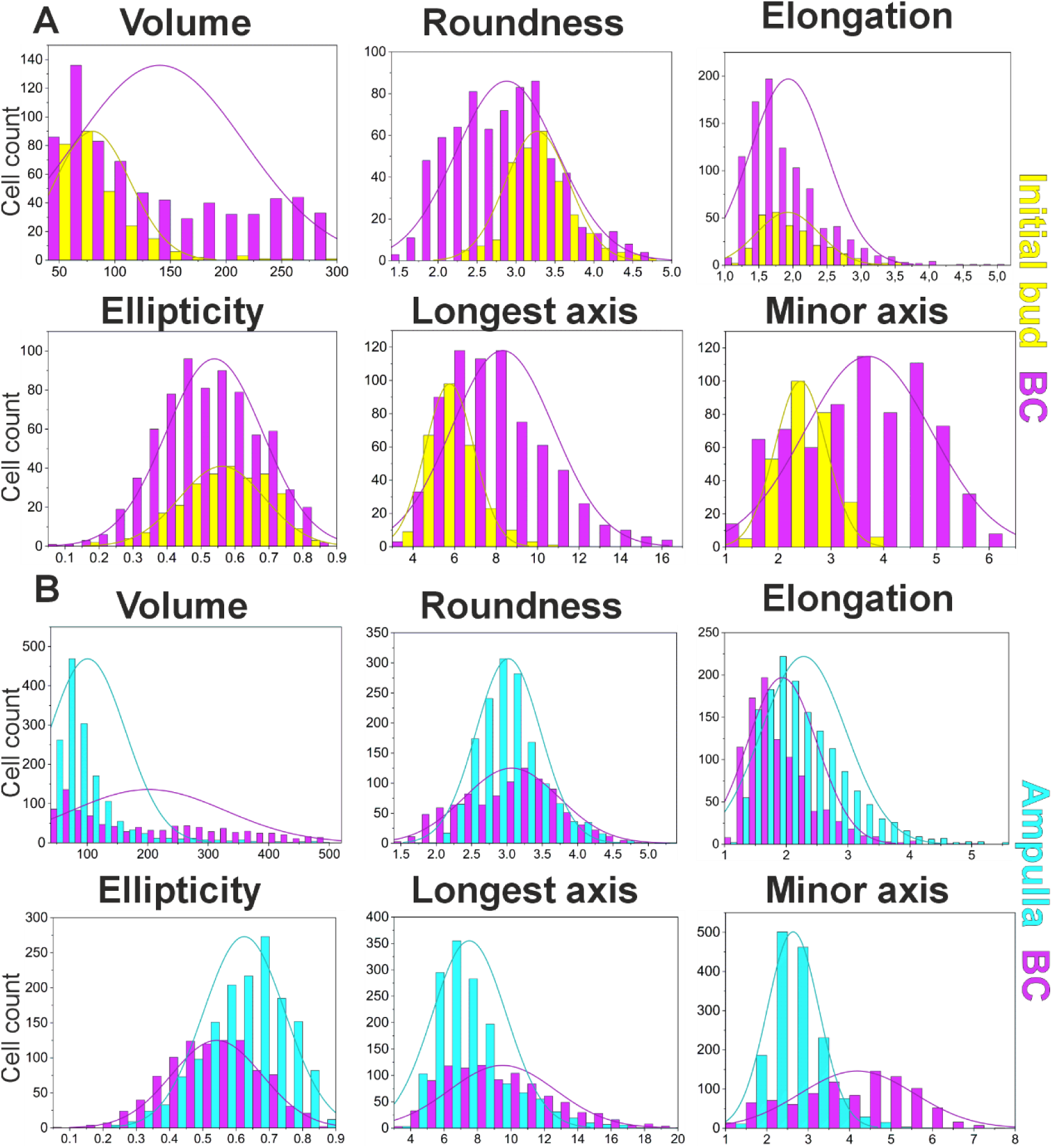
Branching-compromised epithelial cells are in general wide and less elliptical than epithelia in control kidneys. **A)** Distribution plots (cell count on Y axis; microns/cubic microns on X axis) of shared volume range cells (50-297µm^3^) showing range of epithelial cell volumes, roundness, elongation, ellipticity, longest axis, and minor axis in branching-normal (yellow) ureteric bud at initial bud stage. The distribution range of branching-compromised epithelial (BC) cells is shown in violet and demonstrates that these cells have lower values for roundness than those in branching-intact ureteric bud, yet their minor axis is wider than in control cells. **B)** Distribution plots of shared volume range cells (∼50-500µm^3^) showing range of epithelial cell volumes, roundness, elongation, ellipticity, longest axis, and minor axis in branching-intact (turquois) ureteric bud at ampulla stage. Overlay of and comparison to the distribution range in BC epithelial cells (violet) shows that most cells in the control epithelia are small in volume and longest axis while largely of median roundness. BI epithelia show wide distribution of volumes, roundness, and longest axis, while their distribution is skewed towards less elongated and elliptical forms.

Ellipticity and elongation of epithelial cells in BC tips resemble the values in BI epithelium at initial bud stage, but show steady decline when compared to ampulla, asymmetric ampulla, and T-bud stages (**Fig. 7, S4**). The diminished ellipticity and elongation together with high cell volume in BC epithelial cells suggests possible defects to compress cells into elliptical/elongated forms. One mechanism to do this is apical constriction through which cuboidal epithelia acquire typical wedge shape. To test this hypothesis, we measured the longest, intermediate, and minor axes to find out that their median values are higher in BC than in BI cells (Table 1-2). Stage-by-stage comparison of the longest axis distributions identified that it does not deviate between the cells in BI and BC epithelia at asymmetric ampulla and T-bud stages (**Fig. 7, S4**). The BI epithelial cells, however, have higher longest axis at the initial bud stage, and it remains higher also when compared to BI epithelia at the ampulla stage. Notably, the minor axis of BC epithelial cells is consistently higher than in BI cells throughout all the branch cycle stages (**Fig. 7, S4**). These results demonstrate that the same volume range epithelial cells in BC epithelia are on average wider than cells at BI initial bud and ampulla stage. Moreover, BI cells at asymmetrical ampulla and bifurcated tip stages are on average more asymmetrically constricted than UB epithelial cells in BC tips. Taken together, our analyses indicate a failure to shorten minor and intermediate axes, suggesting a defect in the ability of BC epithelium to generate the normal curvature required for the branching process.

### Ureteric bud cells modeling branching incompetence have disrupted lateral adhesions

Our analysis demonstrates that BC epithelial cells are wide and fail to transform into elliptical and elongated shapes. We have previously shown that MAPK/ERK-deficient UB epithelium used here to model branching incompetence has abnormal cell-cell adhesions in the lateral cell borders where intercellular gaps and increased extracellular space are observed.^10^ To further molecularly characterize BC epithelial cells we re-analyzed their transcriptional profiles ^30^ to identify gene expression changes related to cell geometry and adhesion. This revealed that differentially expressed genes in BC epithelium associate with actin polymerization, adherens junctions, apical constriction, focal adhesions, regulation of cell size and shape, and tight junctions (**Fig. S5A**). Also, *Magi3*, which is proposed to maintain steady state intercellular tension within the epithelial sheet,^59^ is downregulated in the BC epithelium.^30^ As suggested by our previous findings,^10^ adherens junctions and focal adhesions had the highest abundance of differential transcripts while several differentially expressed transcripts also fall under the tight junction category, including a UB tip-specific transcript *Wnt11* with a known function in guiding UB branching.^60^

The possible changes in adherens junction dynamics caused by the MAPK/ERK-deficiency and leading to branching defect were next assessed. It has been shown that ERK directly phosphorylates PAXILLIN at serine 83 in medullary collecting duct cells.^61^ Interestingly, our previous findings showed that although pPAXILLIN levels were reduced in the cytosol of MAPK/ERK-deficient UB-derived cells, an increased level of total PAXILLIN, E-CADHERIN (E-CAD) and VINCULIN were detected at the plasma membranes.^10^ To verify this further, an overall 40% increase in surface E-CAD in UB-derived BC epithelial cells was detected by On-Cell Western (**Fig. S5B-C**, P = 0.002996, n =5). As adherence junctions are physically linked to actin cytoskeleton via beta-catenin, we next assessed MAPK/ERK activation in beta-catenin mutant UB epithelium showing abnormal UB branching morphogenesis.^46, 47^ This demonstrated that loss of beta-catenin results in severely reduced pERK1/2 signal while its stabilization via exon 3 deletion increases ERK activation specifically in the UB trunks (**Fig. S5D-F**). These results support the view that MAPK/ERK activity in UB epithelium may participate in sensing biomechanical signals as previously suggested for contractile and extensile forces.^62, 63^

Since the adhesion sites are interdependent of each other, we further assessed whether the focal adhesion changes ^10^ (**Fig. S5A**) are cause or consequence of the disrupted adherence junctions in MAPK/ERK-deficient epithelial cells. UB-derived cells were transfected with green fluorescent protein (GFP)-tagged wildtype PAXILLIN and co-stained with E-cadherin, endogenous paxillin, and overexpressed paxillin. The cell line was also transfected with GFP-tagged, dileucine- and dialanine-modified E-cadherin. PAXILLIN overexpression in UB-derived cells had no effect on E-CAD, whereas excess E-CAD provoked similar mislocalization of PAXILLIN on cell membranes (**Fig. S5G-H**) as detected in our BC UB tips.^10^ Finally, to evaluate the coherence and functionality of the adherence junctions in BC epithelial cells, the adhesive forces between UB cells were quantified by utilizing a modified traction force measurement (TFM) approach with cell doublets (see **Methods** for details). This revealed a significant decrease (*P* = 4.3940e^-6) in cell-cell adhesion force in BC epithelial cells, while only a minor change (*P* = 0.0511) in the traction stresses exerted by the epithelial cell doublets onto the matrices was detected (**Figure 8A-C**).

**Figure 8.**
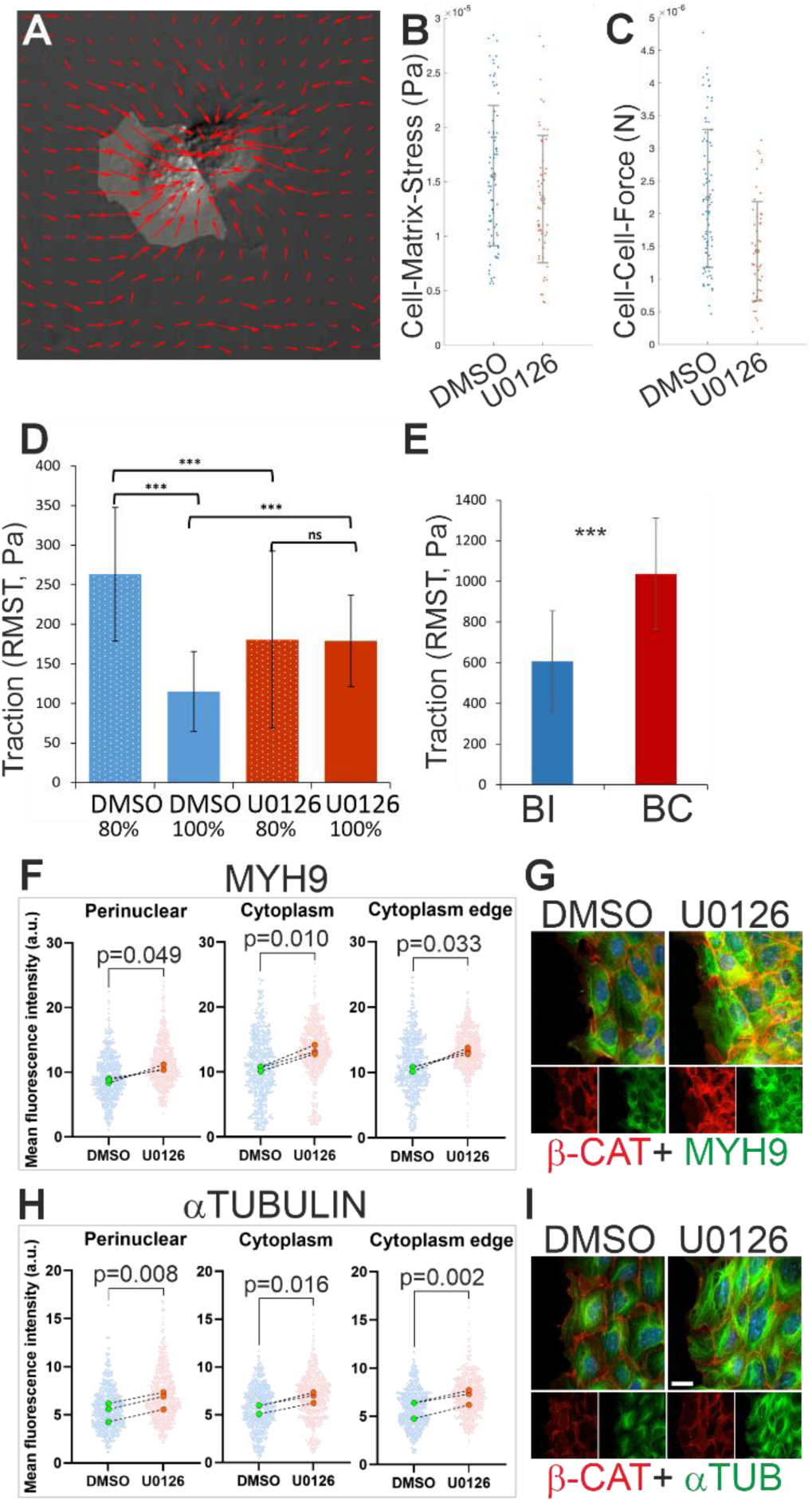
Changes in adhesive forces and actin dynamics contribute to epithelial cell shape modification failure in branching-compromised ureteric buds. **A)** Example of cell doublets and traction generated. **B)** Quantification of stress (Pa) exerted by UB-derived cell doublets treated with U0126/DMSO onto PAA matrices (P = 0.0511). Example force vector map indicating traction forces exerted by UB cell doublets on polyacrylamide gels with Young’s modulus E = 19660 Pa. Measurements were taken from 85 independent doublets across five technical replicates for DMSO control group and from 59 independent doublets across four technical replicates for U0126-treated group. Error bars are +/− standard deviation. Significance was determined by two sample t-test with MATLAB function ttest2. **C)** Quantification of forces (N) between UB-derived cells in doublets treated with U0126 (to mimic branching defect) or DMSO control (*P* = 4.39E^-06). **D)** Average root mean square traction (RMST) forces of UB-derived cell line at ∼80% (*P* = 0.000166) and 100% (*P* = 9.59E-08) confluence on fibronectin-coated 25 kPa gels. 24h U0126 treatment. Example force heatmaps generated from TFM for DMSO and U0126 conditions. **E)** RMST forces in Pa of primary UB cells isolated from 3 control (BI) and 3 UB-specific MAPK/ERK-deficient (BC) embryos at E12 and allowed to delaminate for 48 hours on fibronectin-coated 25 kPa gels before imaging. *P* = 5.69E-06. **F)** Quantification of mean fluorescent intensities of MYOSIN 9 (MYH9) signal in control (DMSO) and MEK-inhibited (U0126) primary ureteric bud (pUB) epithelial cells. **G)** Representative images of MYH9 (green) and BETA-CATENIN (β-CAT, red) co-immunolabeling in control (DMSO) and MEK-inhibited (U0126) pUB epithelial cells. **H)** Quantification of mean fluorescent intensities of alpha-TUBULIN (αTUB) signal in control (DMSO) and MEK-inhibited (U0126) pUB epithelial cells. **I)** Representative images of αTUB (green) and β-CAT (red) co-immunolabeling in control (DMSO) and MEK-inhibited (U0126) pUB epithelial cells. Scale bar: 20µm

### Ureteric bud cells with deficient cell-cell adhesions show compensatory focal adhesion changes

In addition to quantifying cell-cell adhesion at adherens junctions, we also examined the effects of MAPK/ERK-deficiency on focal adhesions using TFM to calculate the cell-matrix traction forces exerted by the cells. Measurements were taken from primary BC UB cells and UB-derived cell line where MAPK/ERK-deficiency was induced by chemical MEK-inhibition for 24 hours. This demonstrated that following MAPK/ERK inactivation by MEK-inhibition, at ∼80% confluence, the UB-derived cell line showed reduction in cell-matrix stress (*P* = 0.000166) (**Figure 8D**). However, at 100% confluence, the UB-derived cell line had increased traction stress towards the matrix upon MAPK/ERK-deficiency (*P* = 9.59E-08). Note that the baseline of root mean square traction (RMST) [Pa] for the control group at 80% confluence is nearly three times the baseline RMST for the same group at 100% confluence (*P* = 2.45E-17). Remarkably, the RMST for the MAPK/ERK-deficient cells was unchanged regardless of confluence (*P* = 0.938587) supporting the findings that the BC epithelial cells are unable to sense biomechanical cues from the surrounding tissues. Notably, BC primary UB cells exerted a significantly higher stress (*P* = 5.69E-06) onto the matrix compared to the wildtype primary UB cells, (**Figure 8E**) suggesting functional deficiency in cell-matrix adhesion.

Cell adhesions are intimately linked to actomyosin that critically anchors cadherins at adherens junctions and together with microtubule network contribute to mechanosensitive regulation of cell shape changes during epithelial morphogenesis.^64, 65^ It has been shown that loss of non-muscle myosin protein functions in UB epithelium causes decreased apical constriction and E-CAD mediated adhesions with simultaneously increased ERK activation.^38^ To test if actomyosin contractility and/or microtubule networks are affected by the loss of ERK activation and thus could contribute to the cell shape changes in our BC UB tips, the localization and intensities of MYH9, α-TUBULIN, and phosphorylated MYOSIN Light Chain (ppMLC) were studied. This showed significantly increased levels of MYH9 and α-TUBULIN in MAPK/ERK-deficient cells (**Fig. S8F-I**), whereas ppMLC levels remained unaffected (**Fig. S6A-B**). The data supports the view that the biomechanical environment in BC UB tips is rigid leading to defects in the ability to non-uniformly compact cellular volume.

### Branching-compromised ureteric bud epithelial cells display disturbed nuclear morphology

Emerging data indicates that changes in cell adhesions are intimately coupled with nuclear morphology and structure.^66, 67^ In this view, the nucleus acts as a dynamic biomechanical sensor of intrinsic and extrinsic mechanical forces according to which it can adapt changes to its shape and function. Since BC epithelial cells demonstrate remarkable changes in their cellular adhesion forces, electron microscopy was utilized to analyze nuclear shapes in wildtype and branching-compromised UB epithelium. This revealed a striking difference in nuclear morphology as nuclei in BC UBs exhibit a highly wrinkled appearance of the nuclear envelope (**Fig. 9A-B**). To quantify these changes, nuclear shape characteristics were measured using Fiji (ImageJ) to find out no change in nuclear size (area or perimeter) of the BC UB (**Fig. S6C-F**). However, there was a significant decrease in the circularity and solidity of the nuclear membranes (**Fig. 9C-D**). The decrease of solidity indicates increased irregularity of the nuclear boundaries (i.e. the nuclear membranes are more wrinkled) while decreased circularity indicates that the nuclei in the BC epithelial cells are more elongated than the nuclei in wildtype UBs.

**Figure 9.**
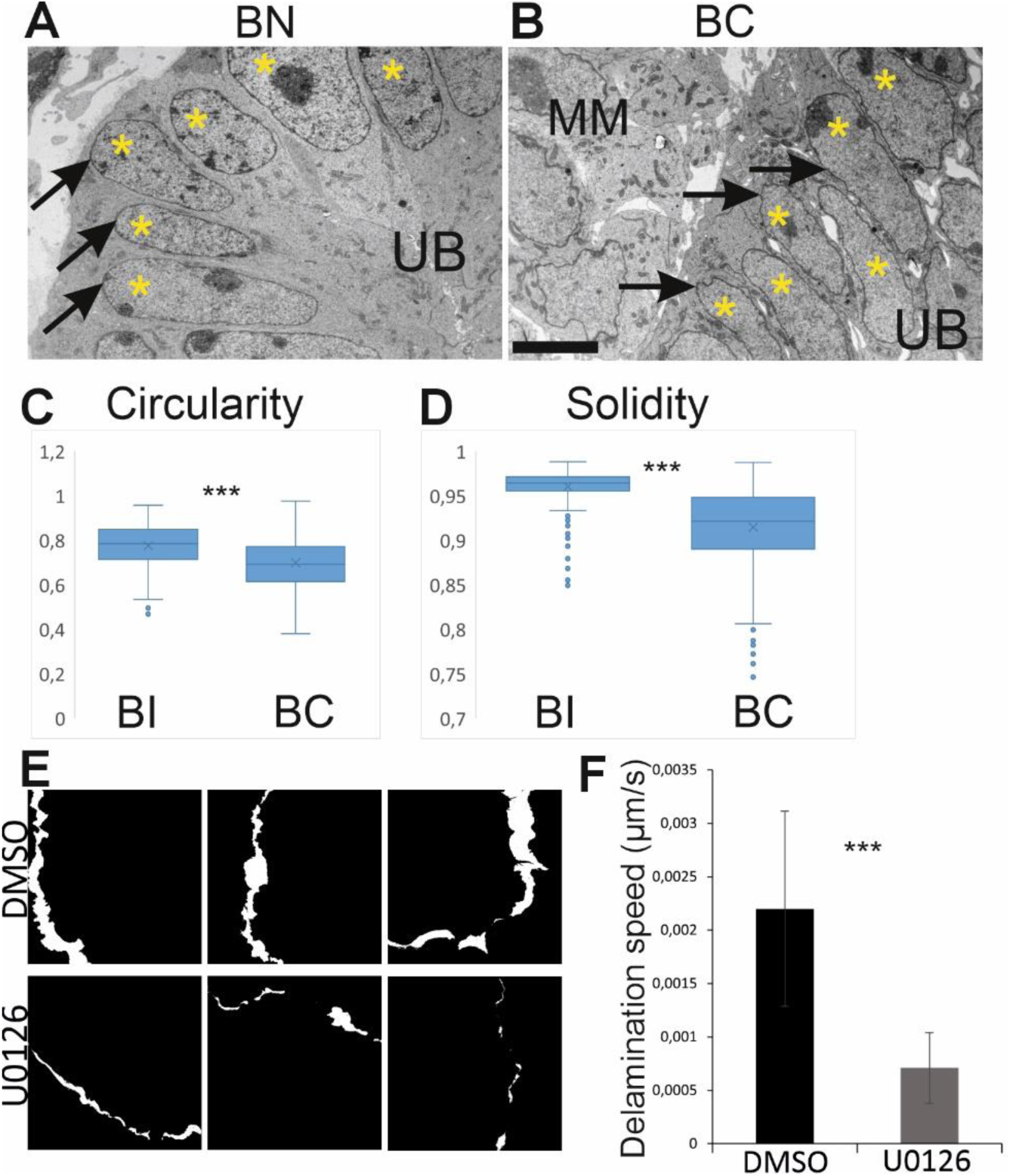
Branching-compromised ureteric bud tips have wrinkled nuclei. **A)** Representative electron microscopy image showing nuclear shapes in branching-intact (BI) ureteric bud epithelium of kidneys at E12.5. **B)** Representative electron microscopy image showing nuclear shapes in branching-compromised (BC) ureteric bud epithelium of kidneys at E12.5. Asterisks decipher ureteric bud epithelial cell nuclei, arrows point to nuclear envelopes, which are wrinkled and irregularly shaped in branching-compromised ureteric buds. **C)** Quantification of nuclear circularity and **D)** solidity (wrinkling) in the branching-intact (BI) and -compromised (BC) ureteric bud epithelial cells show significantly reduced values for BI (*P* < 0.001). **E)** Graphical illustration of leading-edge motilities in control and MAPK/ERK-inhibited primary UB epithelial cells. The result is extracted from live-imaging of actin-reporter (LifeAct)-derived primary UB delamination over 88 minutes (**Supplementary Video 1**) where still images were taken at t=0 min and t=88 mins and subtracted to obtain the resulting masks shown in this figure; white areas were measured to obtain mean distance moved over time. **F)** Quantification of the mean speed of UB delamination (μm/s) measured as the increase in size of delaminating cell masses over 5280 s (88 min) on fibronectin-coated glass. Measurements were taken from 17 independent areas from 5 delaminating primary UBs for DMSO and from 5 independent areas from 2 delaminating UBs for U0126. *P* = 2.14E-05. Error bars are mean +/− standard deviation. Scale bar: 5µm

### Branching-compromised ureteric bud epithelial cells show reduced F-actin rearrangements

The connection of adherens junction proteins to the actin cytoskeleton, the role of actin in maintaining cellular shape and size, as well as the effects of ERK on actin polymerization and leading-edge protrusion during cell motility ^27, 68–72^ altogether led us to assess the dynamics of F-actin in primary cells isolated from UB tips. Primary UBs excised from E11.5 LifeAct-GFP reporter embryos visualizing actin cytoskeleton were plated onto fibronectin-coated, glass-bottom dishes and allowed to delaminate in the control and MAPK/ERK-inhibited conditions. Time-lapse imaging of GFP-labeled actin at the leading edges of the delaminating UBs revealed a clear qualitative defect in actin polymerization and resulting decrease in actin dynamics in MAPK/ERK-deficient cells (**Video S1**). In addition, the distance of expansion of the leading edges of the delaminating cell masses were calculated over the imaging time to give the average delamination speed (**Fig. 9E**). MAPK/ERK-deficient primary UB epithelial cells delaminated significantly slower (*P* = 2.14E-05) than the control cells, at approximately one-fourth of the speed (**Fig. 9F**).

## DISCUSSION

Orchestrated interplay of epithelial branching morphogenesis with the surrounding tissues drives the coordinated differentiation of many mammalian organs. Cellular processes driving epithelial behaviors during tissue deformation include bending, folding, lengthening, and narrowing, which all together modify individual cell shapes and volumes to achieve unique branching strategies in distinct final architecture.^1, 2, 73^ Ureteric bud branching morphogenesis guides growth, patterning, and differentiation of mammalian kidney and while its molecular regulation is rather well established, much less is known about the cellular mechanisms involved in new branch point determination.^3, 74^ Here we utilized artificial intelligence-based ShapeMetrics pipeline to characterize epithelial cell geometries in developing kidney during its different stages of UB branch cycle phases in normal and branching-compromised (BC) kidney, where MAPK/ERK activity is specifically inactivated in UB epithelium.^10, 43^

Our global cell shape analysis identified high heterogeneity throughout all stages of UB branching. Interestingly, heterogeneity in the epithelial cell geometry is high from the very start, already at the initial bud stage, and does not seem to deviate greatly during the later branch cycle stages. Of the individual geometrical parameters, the range of ellipticity and elongation are shared among the different UB morphologies while roundness deviates most similarly as volume, which however shows lesser degree of fluctuation. Interestingly, the number of cells per tip is highest at the ampulla stage suggesting that epithelial crowding precedes the decision on where the new branch site will be located. UMAP projection of the data separated T-bud epithelial cell characteristics from the rest of the stages suggesting that T-bud stage tips analyzed here are not simply just new bud stage UB tips in the reiteration of branching events during kidney development. Geometric parameter-based hierarchical clustering of epithelial cells and their localization studies aiming at understanding the patterns where cells with specific shape profiles land in the UB tips revealed that none of the parameter combinations could be used to predict morphological changes. However, two types of cells displayed tendency to localize to the most cortical regions of the UB tips at asymmetric ampulla and T-bud stage: cells which are simultaneously long, elongated, and elliptical, and cells where all parameters are low. Of these, the most elongated and elliptical cells show patchy pattern predominantly at ampulla and asymmetric ampulla stages, where the next growth directions will be determined. This suggests that cells with such morphologies may represent distinct identities within molecularly uniform UB tip domains of crowed ampullae.

Distinct mammalian cell morphologies reflect their functional specialization and often the control of cell shape is coupled to cell fate decisions.^75^ Here we identified that cell size (volume) is clearly larger in BC UB epithelium than in normally branching UB. Small cell size has been linked to stemness in skin and hematopoietic system, where bigger sized cells show reduced proliferation capacity and increased terminal differentiation.^76–78^ Our previous transcriptomic characterization of BC epithelium revealed a loss of UB tip identity and premature differentiation of collecting duct cell types.^30^ Control of cell shape changes during branching is mediated by an interplay of extrinsic biochemical signals and mechanical forces.^79, 80^ Intrinsic cellular mechanisms mediating the responses of extrinsic stimuli to cell interior include cell adhesions, actomyosin and cytoskeletal rearrangements.^64, 81–84^ Our data suggest that biomechanical signals participate in regulation of UB branching morphogenesis and that MAPK/ERK activity likely mediates mechanical properties of the microenvironment to the cell interior, like previously shown in experiments with human mammary epithelial cells.^62^ Other mechanisms influencing cell shapes and overall epithelial development include extrinsic forces from neighboring cells as well as fluid pressure during epithelial lumen expansion.^85–87^ Adhesion-mediated interactions between cells also contribute to the surface tension energy, which is required for modifying cell geometry and tissue mechanics during morphogenesis and homeostasis.^88, 89^

Analysis of cellular properties in BC UB tips identified clearly less elliptical and elongated epithelial cells than in normally branching kidneys. These epithelial cells fail to decrease their size as demonstrated by the large volumes. They appear also rigid as suggested by reduced actin dynamics and increased levels of MYH9 and αTUB indicating abnormalities in actomyosin and microtubule functions that are involved in cell shape regulation in other branching systems.^65, 73, 84^ Microtubules and actomyosin cytoskeleton resist compressive forces and maintain complex shapes especially in large cells.^90^ Microtubules are controlled through microtubules organizing center, which is directly connected to nuclear lamina and together with actomyosin network aids cell shape regulation.^91, 92^ The nuclei of the BC epithelium appear highly wrinkled supporting defects in cytoskeletal organization and suggesting changes in biomechanical environment. In accordance, we demonstrate here that cellular adhesions show molecular abnormalities and are functionally deficient in BC epithelia, as detected by decreased cell-cell adhesion force and cell-matrix traction stress in UB-derived epithelial cells where MAPK/ERK activity is inhibited. Thus, the inability to generate complex tip shapes in BC UB tips at least partially derives from the stiff cellular niche where normal biomechanical sensing processes are disrupted.

Elegant *in vitro* studies with kidney explants and isolated UB cultures suggest that polarized actin localization and myosin-based apical constriction also influence kidney morphogenesis.^93–95^ Also, a cell autonomous Ras-MAPK signal is required for activation of the actomyosin cytoskeleton to promote apical constriction in the zebrafish posterior lateral line primordium.^96^ Previous studies in developing lung epithelium and salivary gland suggest that apical constriction contributes to the regulation of cell shape changes.^97–99^ Precise control of actomyosin dynamics additionally maintains epithelial integrity and morphology in the mature renal collecting ducts.^35^ Interestingly, previous study reports that epithelial cells round up when their adherens junctions are disrupted.^100^ This may be important for epithelial cell proliferation and motility reported in the UB tips.^41, 42, 57, 101–104^ Thus, the identified changes together with existing data propose a proliferation- and stretching-based model for driving new branch formation in the developing kidney. More specifically, our results suggest that to reach an ampulla stage, cells unevenly increase in volume, partially due to high proliferation,^15^ which triggers crowding at the ampulla stage. Morphogenesis from ampulla to asymmetric ampulla stage requires epithelial cells to stretch and acquire elliptical and elongated shapes (**Fig. 10**). The abnormalities in epithelial cell geometry, adhesion, and cell cytoskeleton likely contribute to the inability of the BC UB epithelium to transform into traditional columnar shape cells, which are typically responsible for curvature formation in epithelia.^105^

**Figure 10.**
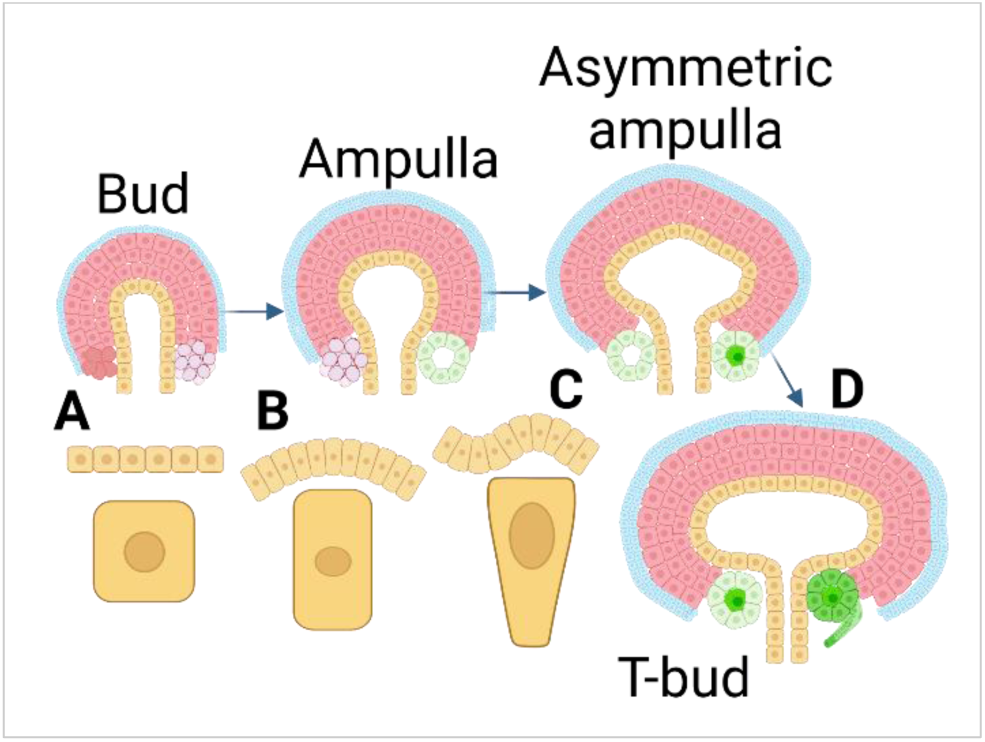
Summary of epithelial morphogenesis during ureteric bud branching in developing kidney. Schematic presentation of an early kidney cell types and main morphological events. Ureteric bud (UB) epithelium is shown in yellow, the surrounding metanephric mesenchymal cells representing nephron and stromal progenitors are depicted in red and blue, respectively. UB epithelium is separately shown below each bud stage to highlight changes in morphologies. **A)** At initial bud stage of UB morphogenesis the epithelial cells are mainly of simple cuboidal type. Subpopulation of nephron progenitors are induced to differentiation as shown by gradually increasing condensation of the cells (lilac). **B)** Epithelial UB tip cells decrease roundness to become more columnar in shape at ampulla stage where more bended epithelial structures appear. Nephrogenesis progresses further as some of the condensates (lilac) transform into pretubular aggregates (light green). **C)** Major morphological changes occur during transition of UB from ampulla-to-asymmetric ampulla morphology as individual epithelial cells unevenly condense their conformation into elliptical and elongated forms to generate local curvatures. Nephron differentiation progresses towards full epithelialization which then at **D)** T-bud stage is shown as comma-shaped bodied nephron precursor (dark green).

## Supporting information

Video_LifeAct_pUB_timelapse

Main tables

Kurtzeborn et al_supplemental files

## Acknowledgements

We thank Agnès Viherä and Maare Arffman for their expert technical skills, Sean Yao for cell-to-matrix traction force measurements, Sara Wickström from advice for cell-cell adhesive force measurements, Santiago Cortés Reina for his MATLAB expertise. We thank Laura Kerosuo for her insights into the interpretation of segmentation results and Peter Hohenstein for comments on manuscript. Imaging was performed at the Light Microscopy Unit (University of Helsinki, Institute of Biotechnology) and the Bio-Imaging Unit (University of Helsinki, Medicum). Part of the work was carried out with the support of HiLIFE Laboratory Animal Center Core Facility, University of Helsinki, Finland and HiLIFE GM-Unit Core Facility, University of Helsinki, Finland; a member of Biocenter Finland. Tissue processing to paraffin was performed at the Tissue Preparation and Histochemistry Unit, Department of Anatomy, University of Helsinki. This work was supported by grants to S.K. from (Aamu Pediatric Cancer Foundation; Research Council of Finland; HiLIFE University of Helsinki; Väre Children’s Cancer Foundation) and to K.K. from Munuaissäätiö and the Orion Research Foundation.

## Author Contributions

K.K., V.I., and S.K. designed the experiments, S.K. conceptualized the project. K.K., V.I., H.A., T.Z., and J.K. conducted the experiments. D.B. provided beta-catenin deficient kidney samples and P.C. performed UMAP analyses. K.K., V.I., and R.K. analyzed data. K.K., V.I., and S.K. wrote and edited the manuscript.

## Data availability

All data needed to evaluate the conclusions in the paper are present in the paper and/or the Supplementary Materials.

