## Supplementary material for "Biomechanical regulation of cell shapes promotes branching morphogenesis of the ureteric bud epithelium": Kurtzeborn et al_supplemental files

**Supplemental Figure 1**.

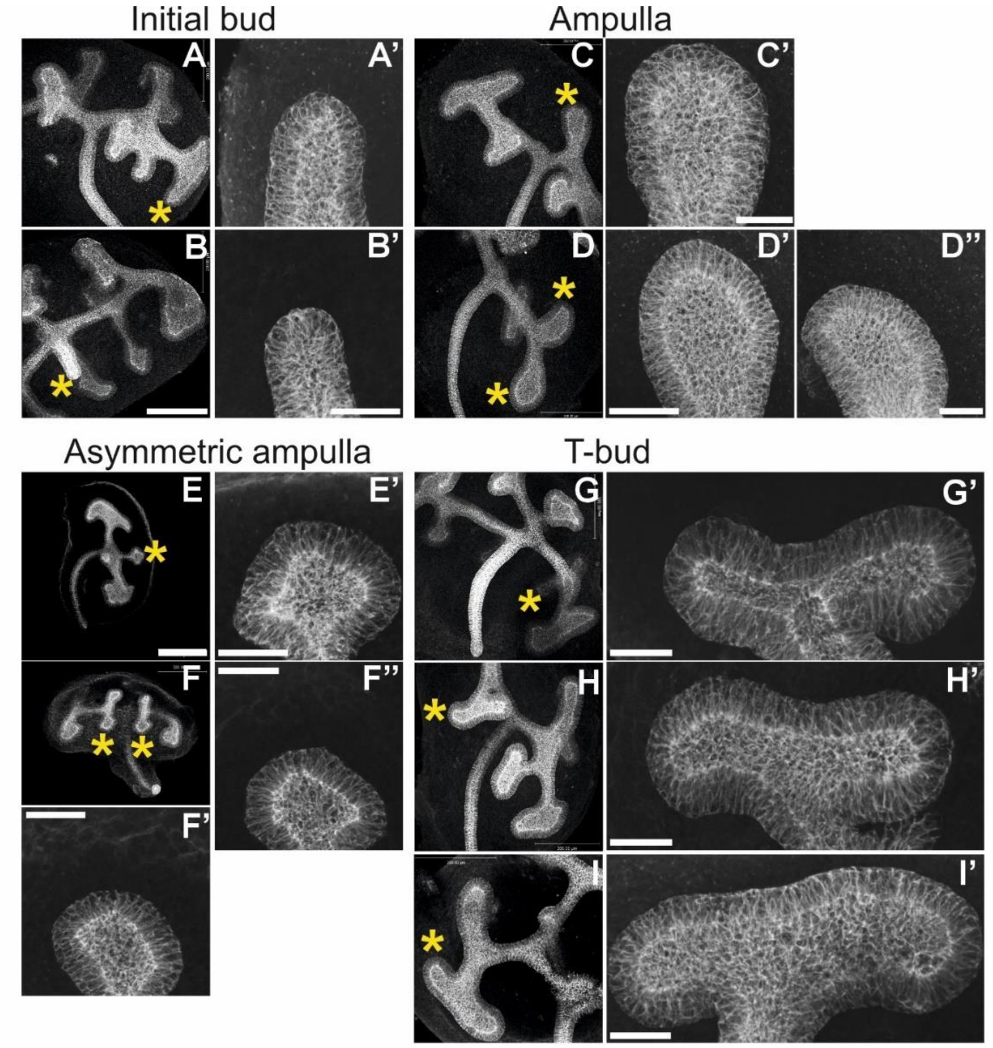


**Supplementary Figure 1**. E12.5 whole kidneys used for ShapeMetrics segmentation analyses. The ureteric bud epithelium was visualized by E-CADHERIN staining (white). **A)** Low magnification image of the kidney from which the first initial bud stage ureteric tip (asterisk) is imaged in 3D with **A')** higher magnification. **B)** Low magnification image of the kidney from which the second initial bud stage ureteric tip (asterisk) is imaged in 3D with **B')** higher magnification. **C)** Low magnification image of the kidney from which the first ampulla stage ureteric bud tip (asterisk) is imaged in 3D with **C')** higher magnification. **D)** Low magnification image of the kidney from which the second and third ampulla stage ureteric bud tips (asterisks) are shown in **D')** and **D'')** with higher magnifications. **E)** Low magnification image of the kidney from which the first asymmetric ampulla stage ureteric bud tip (asterisk) is imaged in 3D with **E')** higher magnification. **F)** Low magnification image of the kidney from which the second and third asymmetric ampulla stage ureteric bud tips (asterisks) are shown in 3D **F')** and **F'')** with higher magnifications. **G)** Low magnification image of the kidney from which the first T-bud stage ureteric bud tip (asterisk) is imaged in 3D with **G')** higher magnification. **H)** Low magnification image of the kidney from which the second T-bud stage ureteric bud tip (asterisk) is imaged in 3D with **H')** higher magnification. **I)** Low magnification image of the kidney from which the third T-bud stage ureteric bud tip (asterisk) is imaged in 3D with **I')** higher magnification. Scale bar for A-I: 200µm, A'-I': 40µm

**Supplementary Figure 2**.

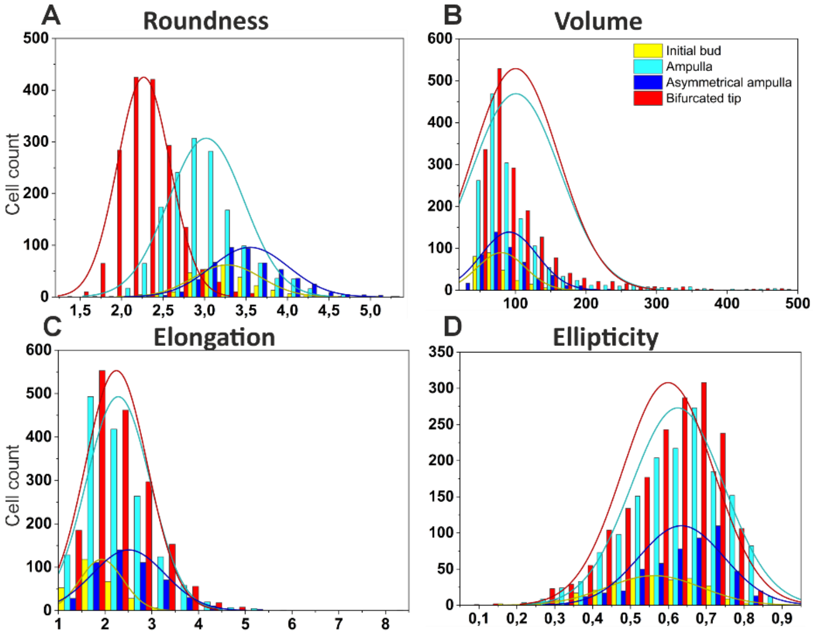


| E | Roundness > 3.2 (%) | Volume > 150 (%) |
|---|---|---|
| Bud | 59 | 3 |
| Amp | 31 | 13 |
| Asym amp | 78 | 7 |
| T-bud | 1 | 13 |

| F | Elongation > 0.23 (%) | Ellipticity > 0.7 (%) |
|---|---|---|
| Bud | 17 | 18 |
| Amp | 41 | 29 |
| Asym amp | 36 | 34 |
| T-bud | 66 | 21 |

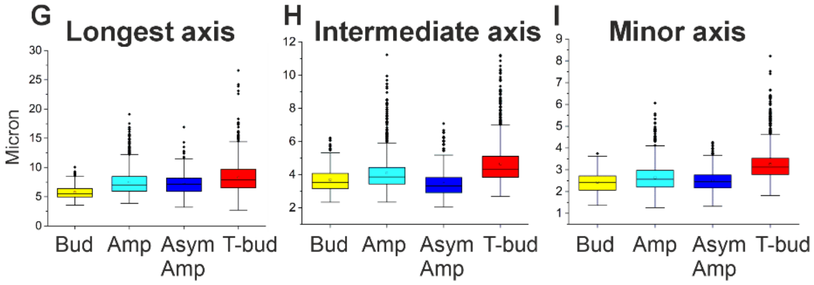


**Supplemental figure 2.** Overlay of distribution plots within the whole volume range (cell count on Y axis; microns/cubic microns on X axis) showing changes in **A)** roundness, **B)** volume, **C)** elongation, and **D)** ellipticity of ureteric bud epithelial cells across the different branching cycle stages. **E)** Summary of representation (percentage) of ureteric bud epithelial cells with higher than 3.2 roundness and larger than 150 $\mu m^2$ volume during the progression of ureteric bud branch cycle. **F)** Summary of representation (percentage) of epithelial cells with elongation higher than 0.23 and ellipticity greater than 0.7 during the progression of ureteric bud branch cycle. Quantification of **G)**

longest, **H)** intermediate, and **I)** minor axis in ureteric bud epithelial cells across different stages of ureteric bud branch cycle. Abbreviations: Bud; initial bud, Amp; ampulla stage, As amp; asymmetric ampulla, T-bud; t-bud stage.

**Supplementary Figure 3**.

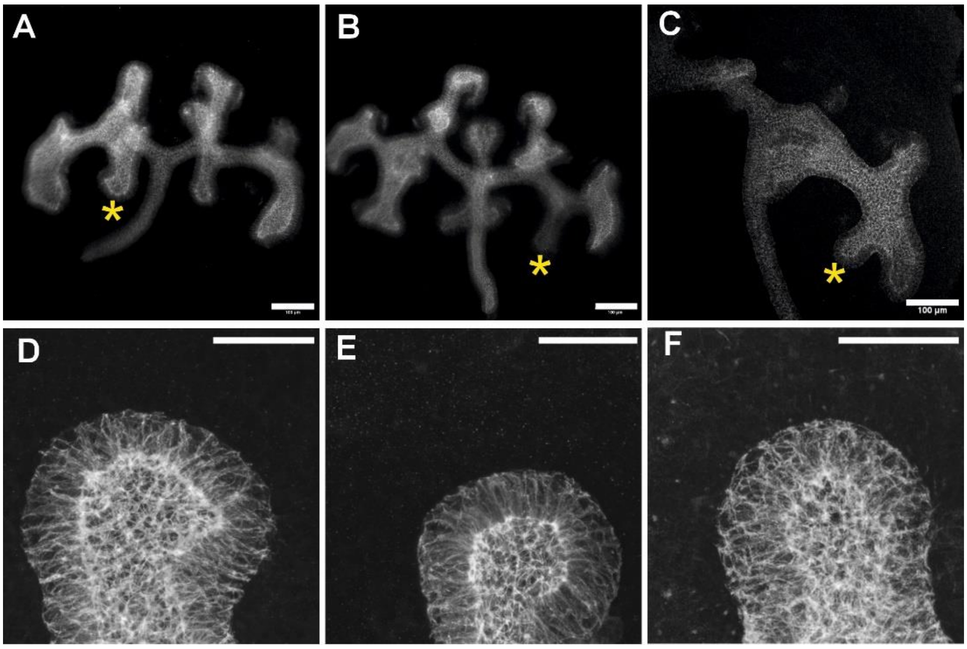


**Supplementary Figure 3.** Branching-compromised E12.5 whole kidneys were derived from genetic loss-of-function model of MAPK/ERK activity specific to the ureteric bud epithelium (*HoxB7CreGFP;Mek1*$^{fl/fl}$;*Mek2*$^{-/-}$). The ureteric bud epithelium was visualized by E-CADHERIN staining (white). **A-C)** Three distinct branching-compromised kidneys used for ShapeMetrics segmentation analyses. **D)** High magnification image of branching-compromised ureteric bud tip denoted by asterisk in A. **E)** High magnification image of branching-compromised ureteric bud tip denoted by asterisk in B. **F)** High magnification image of branching-compromised ureteric bud tip denoted by asterisk in C. Scale bar A-C: 100µm, D-I: 40µm.

**Supplementary Figure 4.**

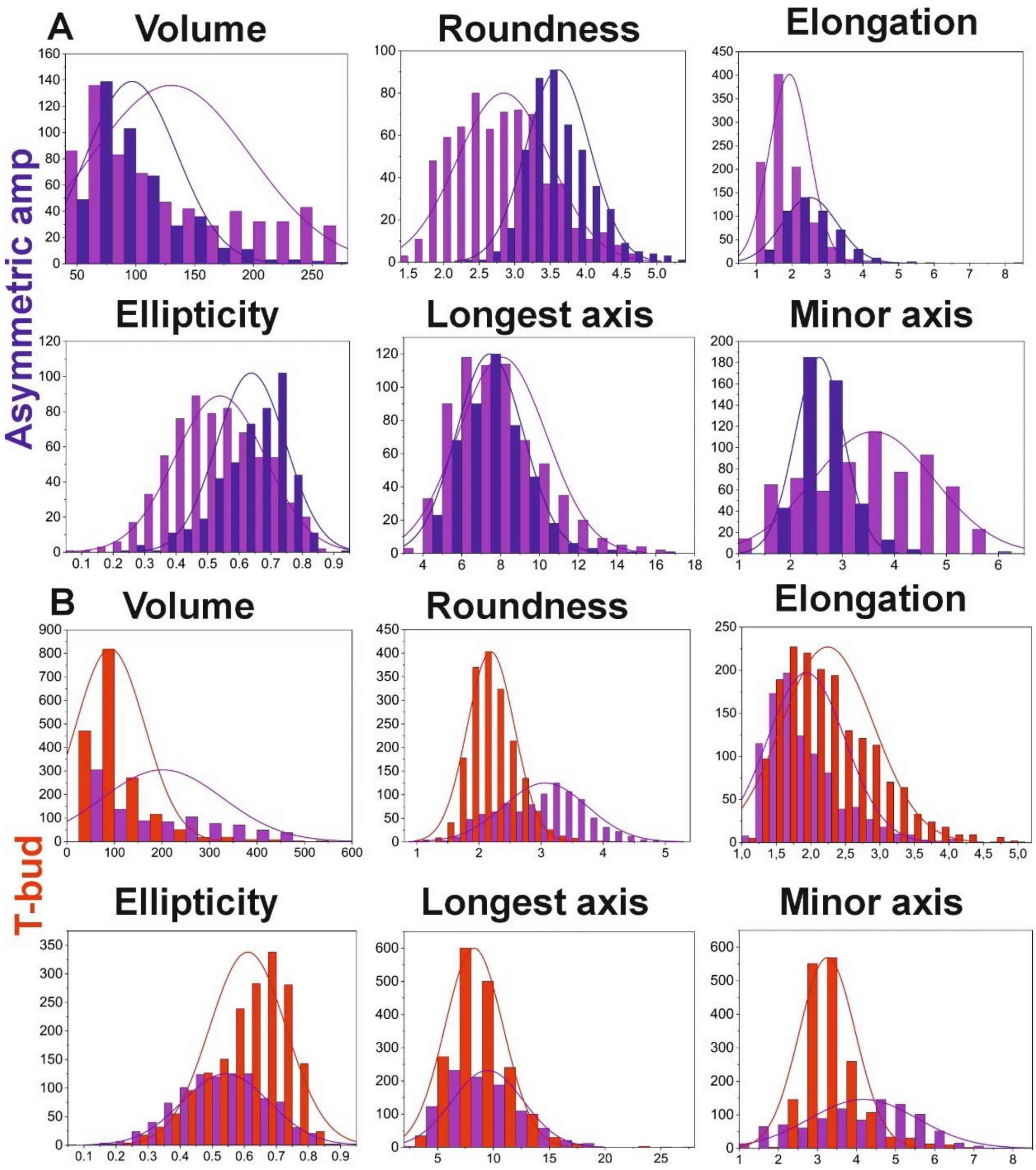


**Supplementary Figure 4. A)** Distribution plots (cell count on Y axis; microns/cubic microns on X axis) showing comparison of epithelial cell volumes, roundness, elongation, ellipticity, longest axis, and minor axis in branching-intact (blue) and -compromised (violet) ureteric buds at the asymmetric ampulla stage. Distributions were compared for cells withing the shared volume range 50-272µm$^3$. **B)** Distribution plots showing comparison of epithelial cell volumes, roundness, elongation, ellipticity, longest axis, and minor axis in branching-intact (red) and compromised (violet) ureteric bud at T-bud stage of ureteric bud branch cycle. Cell volume range in branching-compromised tips and T-bud tips was inherently the same ~50-500µm$^3$.

**Supplementary Figure 5.**

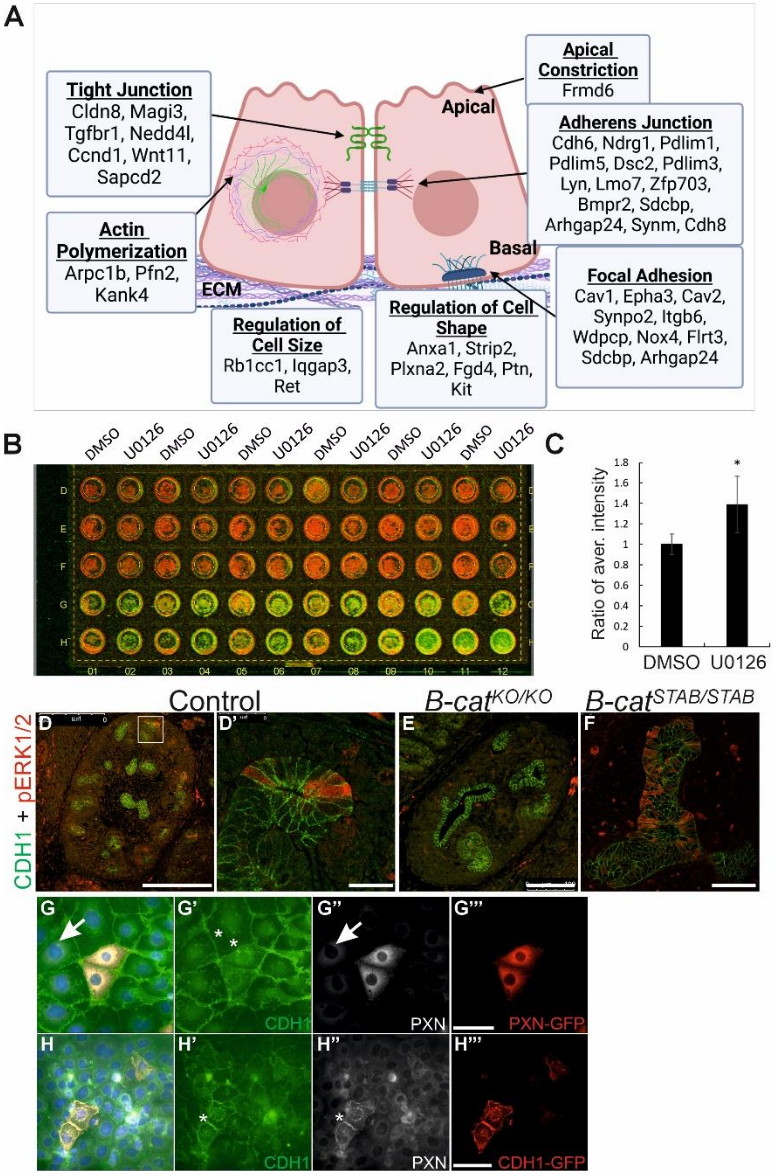


**Supplementary Figure 5. A)** Graphical illustration of differentially expressed genes (DEGs) in branching-compromised ureteric bud epithelium. The cellular localization of DEGs and association with processes related to adhesion and size/shape regulation is shown by arrows. **B)** Representative image of On-Cell Western results showing fluorescently detected E-CADHERIN on ureteric bud-derived epithelial cells. Every other column is control (DMSO) and every other treated with MEK inhibitor (U0126). Three upper rows (D, E, F) are negative controls without primary-antibody while the two bottom rows show immunoreactivity of anti-mouse E-CADHERIN. **C)** Quantification of E-CADHERIN intensity as ratio of intensity average (Y axis). E-CADHERIN intensity is significantly lower in MEK-inhibited epithelial cells than in control cells ($P < 0.05$).  Localization of E-CADHERIN (CDH1, green) and phospho-ERK1/2 (pERK1/2, red) in in E12.5 **D)** Hoxb7Cre control, **E)** tissue-specific knockout (KO) of Beta-catenin (B-cat), and **F)** Beta-catenin stabilized (stab) kidneys reveals loss of

MAPK/ERK activation in the absence of *B-cat* and premature, increased signal when *B-cat* is stablized. **G-G''')** Representative images of PAXILLIN overexpression in ureteric bud epithelial cell line at 48h timepoint. **G)** An overlay of staining with E-CADHERIN (CHD1, green), PAXILLIN (PXN, white) and green fluorescent protein (red) tagged to overexpressed PAXILLIN. Arrow points to endogenous PAXILLIN expression. **G')** E-CADHERIN staining shows similar levels of protein in both wild type and PAXILLIN overexpressing cells (asterisks). **G'')** Immunofluorescent detection and visualization of PAXILLIN only demonstrates both endogenous (arrows) and overexpressed protein. **G''')** Visualization of overexpressed PAXILLIN only by GFP-antibody staining. **H-H''')** Representative images of E-CADHERIN overexpression in ureteric bud epithelial cell line at 24h timepoint. **H)** An overlay of staining with E-CADHERIN (CHD1, green), PAXILLIN (PXN, white) and green fluorescent protein (red) tagged to overexpressed E-CADHERIN. **H')** Immunofluorescent detection and visualization of E-CADHERIN only shows increased protein in cells with overexpression (asterisk) and normal levels in the cells without overexpression. **H'')** Visualization of PAXILLIN only demonstrates intensified protein localization in the epithelial cells overexpressing E-CADHERIN (asterisk). **H''')** Visualization of overexpressed E-CADHERIN only by GFP-antibody staining. Scale bar: D, 250µm; D', F, 25µm; E, 100µm, G-H 50µm

**Supplementary Figure 6.**

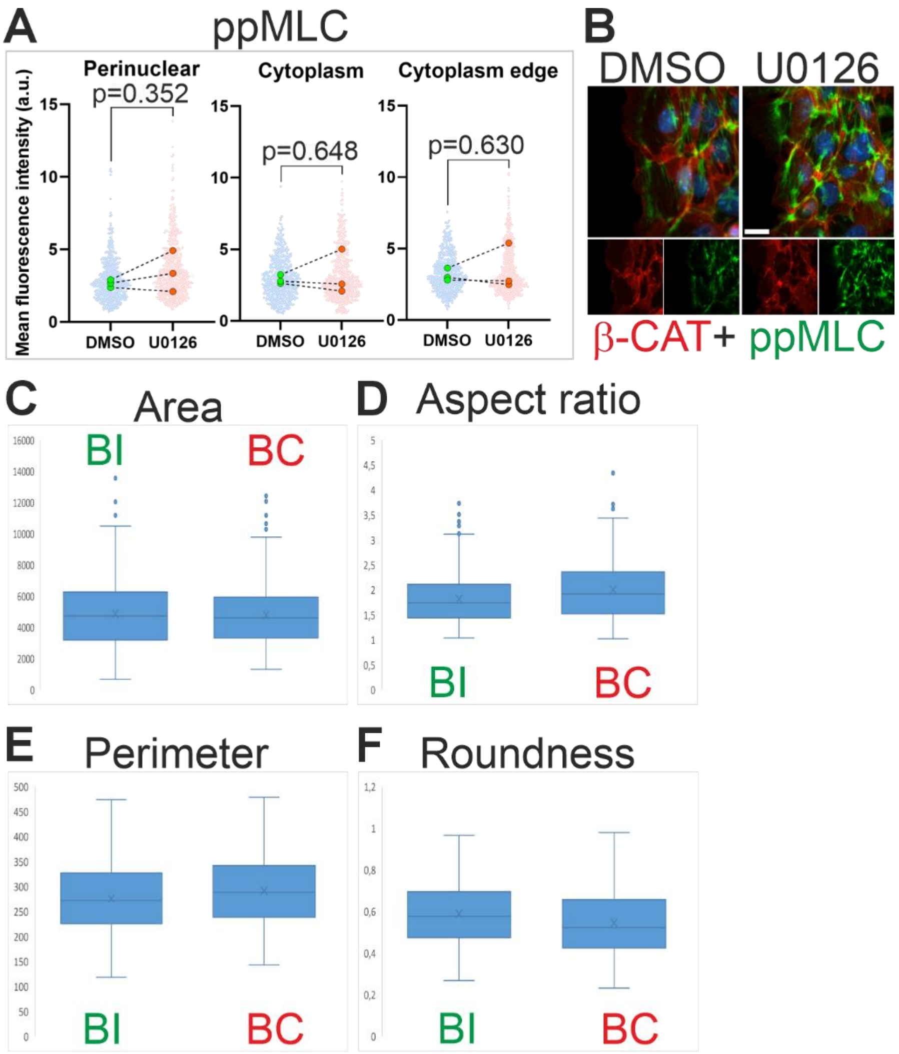


**Supplementary Figure 6. A)** Quantification of mean fluorescent intensities of phosphorylated MYOSIN Light Chain (ppMLC) signal in control (DMSO) and MEK-inhibited (U0126) pUB epithelial cells. **B)** Representative images of ppMLC (green) and β-CAT (red) co-immunolabeling in control (DMSO) and MEK-inhibited (U0126) pUB epithelial cells. The antibody staining signal was measured in distinct subcellular compartments (perinuclear, cytoplasm in the middle and cytoplasm edge at the leading edge) and shown separately in each graph (E, G, I). Quantification of nuclear **C)** area, **D)** aspect ratio, **E)** perimeter, and **F)** roundness in the electron microscopy images of branching-intact (BI) and –compromised (BC) ureteric bud epithelial cells. Scale bar: 20µm
